# Single-molecule optical mapping enables quantitative measurement of D4Z4 repeats in facioscapulohumeral muscular dystrophy (FSHD)

**DOI:** 10.1101/286104

**Authors:** Yi Dai, Pidong Li, Zhiqiang Wang, Fan Liang, Fan Yang, Li Fang, Yu Huang, Shangzhi Huang, Jiapeng Zhou, Depeng Wang, Liying Cui, Kai Wang

## Abstract

**Purpose:** Facioscapulohumeral Muscular Dystrophy (FSHD) is a common adult muscular dystrophy. Over 95% of FSHD cases are associated with contraction of the D4Z4 tandem repeat (~3.3kb per unit) at 4q35 with a specific genomic configuration (haplotype) called 4qA. Molecular diagnosis of FSHD typically requires pulsed-field gel electrophoresis with Southern blotting. We aim to develop novel genomic and computational methods for characterizing D4Z4 repeat numbers in FSHD.

**Methods:** We leveraged a single-molecule optical mapping platform that maps locations of restriction enzyme sites on high molecular weight (>150kb) DNA molecules. We developed bioinformatics methods to address several challenges, including the differentiation of 4qA with 4qB alleles, the differentiation of 4q35 and 10q26 segmental duplications, the quantification of repeat numbers with different enzymes that may or may not have recognition sites within D4Z4 repeats. We evaluated the method on 25 human subjects (13 patients, 3 individual control subjects, 9 control subjects from 3 families) labeled by the Nb.BssSI and/or Nt.BspQI enzymes.

**Results:** We demonstrated that the method gave a direct quantitative measurement of repeat numbers on D4Z4 repeats with 4qA allelic configuration and the levels of post-zygotic mosaicism. Our method had high concordance with Southern blots from several cohorts on two platforms (Bionano Saphyr and Bionano Irys), but with improved quantification of repeat numbers.

**Conclusion:** While the study is limited by small sample size, our results demonstrated that single-molecule optical mapping is a viable approach for more refined analysis on genotype-phenotype relationships in FSHD, especially when post-zygotic mosaicism is present.

## BACKGROUND

Facioscapulohumeral Muscular Dystrophy (FSHD) is a genetic disorder mainly affecting skeletal muscle. The disease progresses in a distinctive pattern and distribution. Clinical symptoms usually appear during the second decade. Weakness begins in the face, shoulders and upper arms, then followed by distal lower extremities, pelvic girdle and abdominal muscles. The symptoms often show marked asymmetry [1]. Several other extra-muscular manifestations are also frequently observed in FSHD, with a high frequency of hearing loss (~75% patients) and retinal telangiectasia (60% patients) [2], as well as defects in the central nervous system such as severe intellectual disability and epilepsy [3]. FSHD is the third most common form of muscular dystrophy and affects as high as 1 in 8,500 individuals in the Netherlands [4]. A significant variability in clinical expression is often observed, even among affected family members.

Two genetic subtypes of FSHD have been identified. The classical form, FSHD1, accounts for 95% of patients and is associated with a polymorphic macrosatellite repeat array on chromosome 4q35. The ~3.3kb repeat unit is referred to as D4Z4, and patients typically carry 1-10 repeats, whereas non-affected individuals possess 11-150 repeats. Another D4Z4 repeat array on chromosome 10q26 exhibits almost complete sequence identity (~99%) to the 4q35 array [5, 6]. Each D4Z4 repeat unit has a complex sequence structure, with several GC-rich repeat sequences and an open reading frame containing two homeobox sequences designated as *DUX4* (double homeobox 4) [7, 8]. Both the 4q35 and 10q26 D4Z4 repeat arrays are highly polymorphic and exhibit extensive size variation in the normal population, but only the 4q35 repeats are associated with FSHD. A polymorphic segment of 10 kb directly distal to D4Z4 on 4q35 was identified [9], with two alleles as 4qA and 4qB, and only the 4qA allele is pathogenic for FSHD [10]. *DUX4* transcripts within D4Z4 are efficiently polyadenylated and are more stable when expressed from 4qA background, suggesting that FSHD1 arises through a toxic gain of function attributable to the stabilized distal *DUX4* transcript [11]. A less common form, FSHD2, accounts for 5% of patients and is associated with the *SMCHD1* (Structural Maintenance of Chromosomes Hinge Domain 1) gene on chromosome 18p11.32 [12]. In FSHD2, patients harboring mutation in *SMCHD1* have a profound hypomethylation of chromosomes 4 and 10, allowing chromosome 4 to express the toxic *DUX4* transcript. Aberrant *DUX4* expression triggers a deregulation cascade inhibiting muscle differentiation, sensitizing cells to oxidative stress, and inducing immune responses and muscle atrophy [13]. A unifying pathogenic hypothesis for FSHD emerged with the recognition that the FSHD-permissive 4qA haplotype corresponds to a polyadenylation signal that stabilizes the *DUX4* mRNA, allowing the toxic protein DUX4 to be expressed [13].

Molecular diagnosis of FSHD is complicated by the relatively large size (~3.3kb) and variable number of repeat units, the presence of homologous polymorphic repeat arrays on both chromosomes 4 and 10, as well as possible exchanges between these chromosomal regions. A Southern blot-based method, a BglII-BlnI dosage test, was developed to improve the sensitivity of conventional Southern blot for molecular diagnosis of FSHD [14]. The method was improved later using restriction enzyme Xap1 which complements Bln1, as the former uniquely digests repeat units derived from chromosome 4 and the latter uniquely digests those derived from chromosome 10 [15]. After these developments, Southern blot could be used as a successful molecular diagnosis test of FSHD. However, the limitations of this method are evident: Southern blot needs four separate time-consuming enzyme/probe combinations and is semi-quantitative, since it estimates repeat count by size of a band in the gel and cannot assay very long alleles. Several attempts have been made to develop alternative methods for diagnosis of FSHD to overcome complications of Southern blot analysis, including molecular combing [16, 17], PacBio sequencing [18], and Nanopore sequencing [19]. Molecular combing has been used in the clinical context, and long-read sequencing in different platforms is also shown to be capable of determining the macrosatellite repeat number since they can generate reads that span over 100kb regions.

In the current study, we aim to leverage a single-molecule optical mapping approach to characterize D4Z4 repeats in FSHD due to repeat contraction. This technique, also referred to as next-generation mapping, has been used for detection of structural variants [20], and Barseghyan et al are among the first to use it in clinical settings, to identify pathogenic SV in a series of patients diagnosed with Duchenne muscular dystrophy (DMD) [21]. Unlike DMD, several key challenges in our study are the differentiation of 4qA alleles with 4qB alleles, the differentiation of 4q35 and 10q26 regions, the accurate quantification of repeat numbers, and the ability to detect low levels of post-zygotic mosaicism.

## RESULTS

### Optical mapping on long DNA molecules can differentiate paralogous genomic regions

Optical mapping [22] is a technique for mapping locations of restriction enzyme sites in DNA molecules. In the current study, we applied a high-throughput platform, the Bionano Saphyr Genome Mapping platform, to perform optical mapping on high molecular weight (>150kb) DNA molecules. The presence of highly similar D4Z4 repeat units in both 10q26 and 4q35 chromosome regions complicates the genome mapping of DNA molecules originating from these regions. To illustrate this, we plotted the pair of segmental duplications that contain the D4Z4 repeat units (DUX gene clusters) (Figure 1A, 1B). Only the cluster in 4q35 are related to the development of FSHD, when there is a contraction of copy number of the D4Z4 repeat units in this cluster. Furthermore, a polymorphic segment of 10 kb directly distal to D4Z4 on 4q35 carry two different allelic configurations in the population, commonly referred to as 4qA and 4qB, and only the 4qA allele is pathogenic for FSHD [10]. An additional variant of 4qA allele is the 4qA-L allele, with a slightly longer D4Z4 unit at the most distal end.

**Figure 1:**
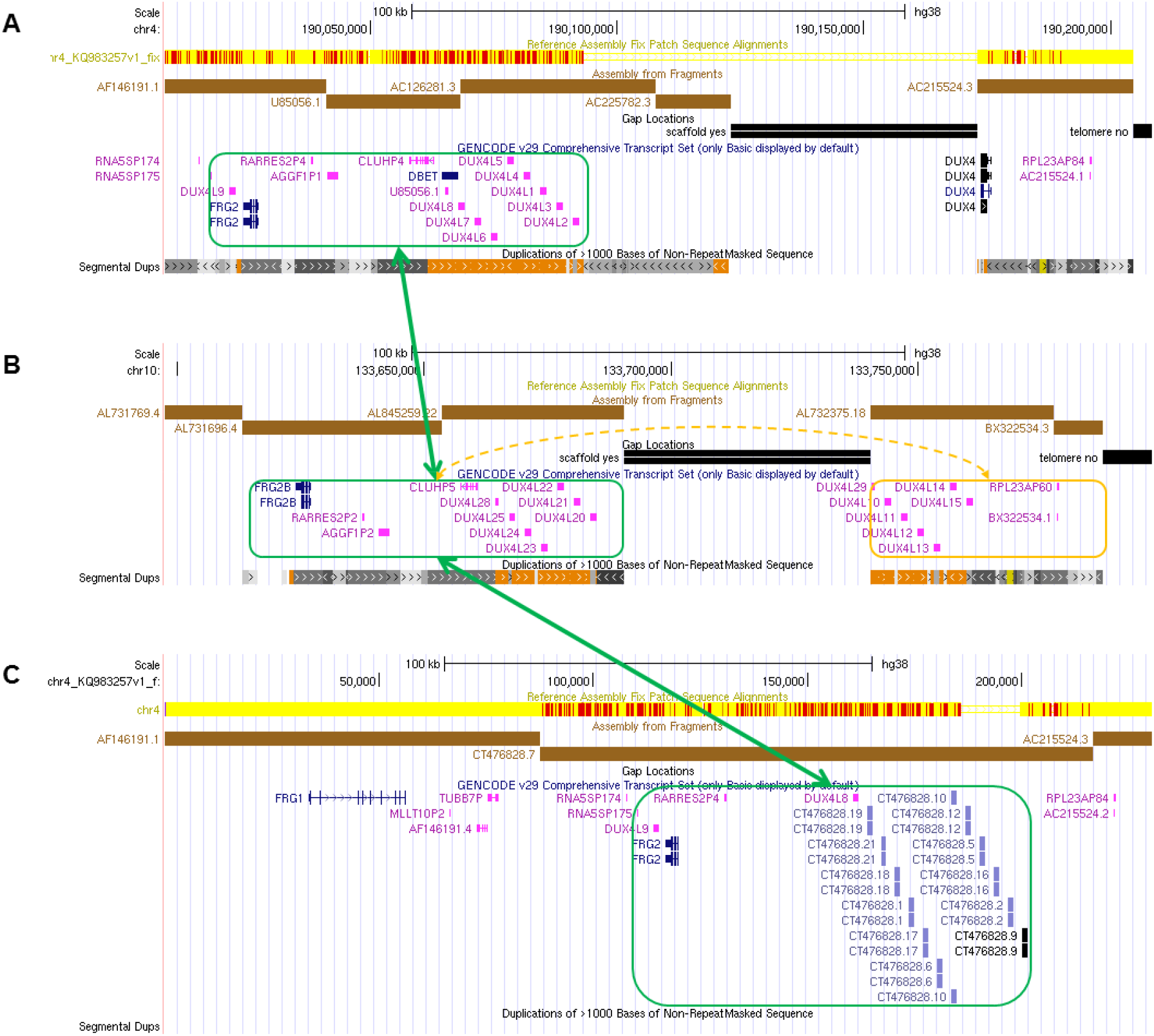
An overview of the genomic architecture of segmental duplications at the chromosome 4q35 (panel A) region, the 10q26 (panel B) region and the KQ983257.1 patch scaffold (panel C). In the GRCh38 reference genome, 4q35 carries one D4Z4 array (8 D4Z4 repeat units); however, 10q26 carries two D4Z4 arrays (each with 7 D4Z4 repeat units), with an incorrect 50kb gap between AC225782.3 (4qB) and AC215524.3 (4qA). A new patch scaffold sequence was recently added in CRCh38 patch 7 (KQ983257.1) with 4qA configuration without gap, and we additionally illustrated the segmental duplication in this patch scaffold. The segmental duplication (green boxes) in 4q35 has high sequence identity with the corresponding region in 10q26 and KQ983257.1, while the distal D4Z4 array separated by the gap in 10q26 is marked with an orange box.

In the GRCh38 reference genome, 4q35 carries one D4Z4 array with 4qB configuration (8 D4Z4 repeat units); however, due to the presence of an incorrect 50kb gap in genome assembly, 10q26 carries two D4Z4 arrays (each with 7 D4Z4 repeat units), while this should be a single continuous D4Z4 array on both chromosomes. We note that GRCh37 does not contain this mistake and shows the correct chromosome 4 and 10 assemblies, but it has a 4qB configuration in chromosome 4. Recently, a new scaffold covering this region was incorporated into GRCh38 patch 7 (KQ983257.1) without the gap, and we additionally illustrated the segmental duplication in this 230kb scaffold, which has a 4qA configuration with 13 D4Z4 repeats (Figure 1C). In summary, successful molecular diagnosis of FSHD requires accurate quantification of D4Z4 repeats within the 4q35 region with high specificity, and the differentiation of the 4qA and 4qB alleles.

### Determining D4Z4 repeat copy number on 4q35 by Nb.BssSI enzyme labeling

We initially selected the Nb.BssSI restriction enzyme for the study, since each D4Z4 unit contains a recognition site for this enzyme, as previously suggested [23]. Through the analysis of fluorescence labels generated by *in silico* digestions with the Nb.BssSI enzyme, we have identified the regions of label similarity and dissimilarity between the paralogous genomic regions on chromosome 4q35 and 10q26 (Figure 2). Our computational analysis showed that when a DNA molecule is long enough to stretch out into the region of dissimilarity (Figure 2, blue box), we can determine whether the molecule originates from 4q35 versus 10q26. Furthermore, when using Nb.BssSI enzyme, based on the presence or absence of one additional label ~1.8kb distal to the last label in the D4Z4 repeat regions, we can determine the allelic configuration to be 4qA or 4qB.

**Figure 2.**
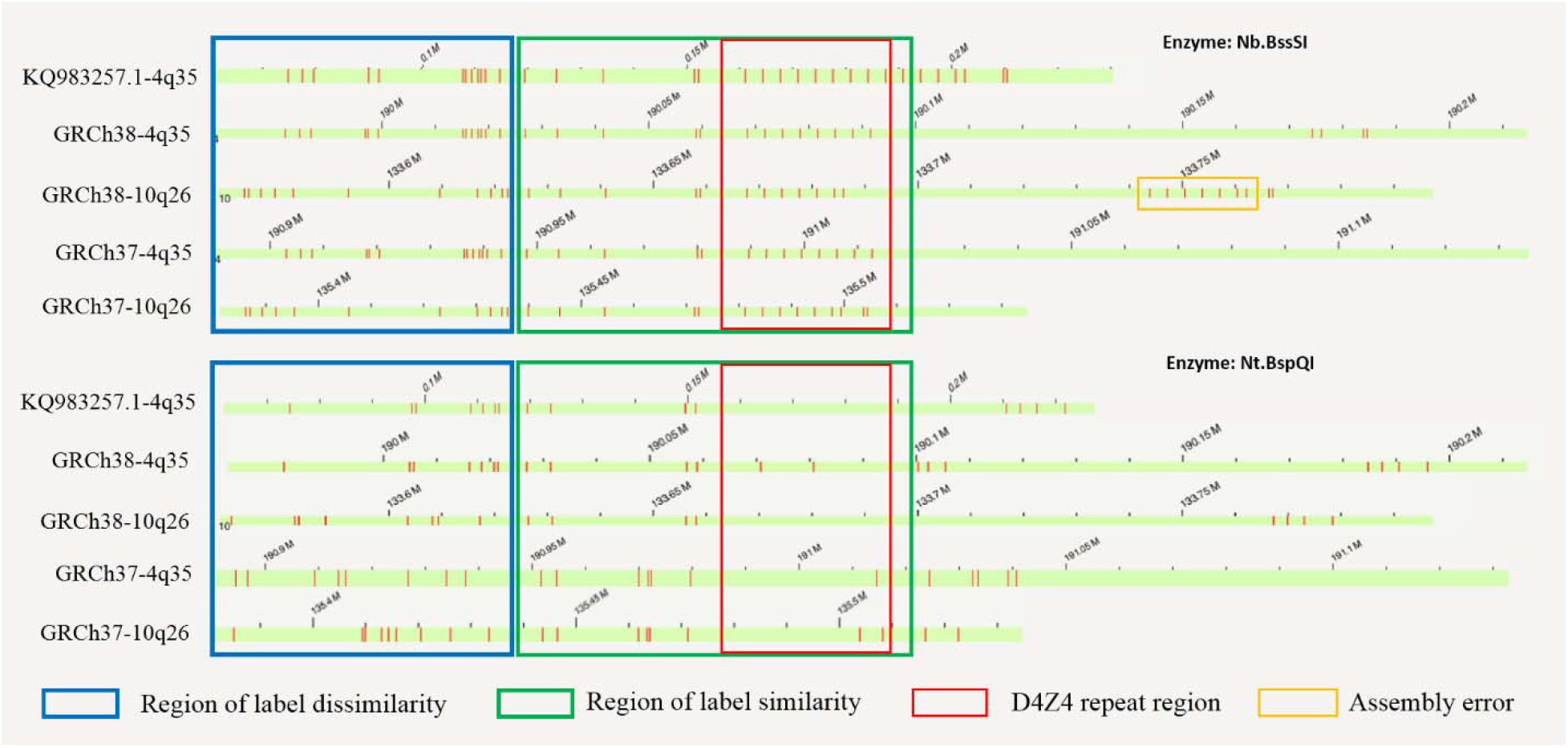
Illustration of the region with label dissimilarity between 10q26, 4q35 and KQ983257.1 (blue box) adjacent to the region of similarity (green box), based on in silico analysis on GRCh37, GRCh38 and KQ983257.1. By using fragments that spans the region of dissimilarity, we can confidently separate fragments originating from 10q26, 4q35 or those that are undetermined (uninformative). The upper and lower panel represent labels generated by the Nb.BssSI and Nt.BspQI enzymes, respectively. Although the reference genome GRCh38 contains two labels (repeat unit #2 and #5) within the D4Z4 repeat region for the Nt.BspQI enzyme (red box), we rarely observe them in real data, possibly due to the inclusion of a very rare allele in the GRCh38 or due to errors in genome assembly. Red vertical bars represent labels of enzyme recognition sites.

We evaluated this approach on five patients who were well characterized clinically (Table 1, Supplementary Table 1). All of them had a positive diagnosis of FSHD through Southern blot (Figure 3), and two of them (ID: P04 and P05) were known to carry post-zygotic mosaicism (as denoted by the “+” sign in Figure 3). The analysis of mapped reads confirmed the presence of contracted alleles in all five patients (Table 1, Supplementary Table 2). We illustrate the observed label patterns on all patients in **Supplementary Figure 1**, where the presence of fluorescence labels can be easily and directly visualized and counted in the Bionano Access browser. Although some enzymatic labels appear to be missing given the imperfect labeling efficiency of the enzyme, it is possible to computationally predict the missing labels based on distances between neighboring labels representing D4Z4 repeats (see Methods). When multiple reads are piled at the same locus, each read will be assigned an estimated repeat count, and a peak-finding algorithm was used to identify the presence of peaks and yield an estimate of the repeat counts.

**Table 1:** A list of patients and control subjects assayed by the Bionano Saphyr platform in the current study. More detailed description of results can be found in Supplementary Table 1. EMG: Electromyography, CK: creatine kinase.

|  | ID | Sex | Age<br>(years) | Onset<br>(years) | Family<br>history | CK<br>(U/L) | EMG | Southern Blot (4q35) |  |  | Optical mapping (Nb.BssSI<br>enzyme, 4q35) |  |  | Optical mapping (Nt.BspQI<br>enzyme, 4q35) |  |  |
| --- | --- | --- | --- | --- | --- | --- | --- | --- | --- | --- | --- | --- | --- | --- | --- | --- |
|  |  |  |  |  |  |  |  | Length<br>( Kb ) | Units | Allele | Units | Allele | Read<br>Count | Units | Allele | Read<br>Count |
| Patient cohort 1 | P01 | F | 37 | 28 | + | 104 | mild<br>myopathic<br>change | ~20 | ~4 | 4qA | 4 | 4qA | 6 | 4.3±0.3 | 4qA | 38 |
|  |  |  |  |  |  |  |  | ~38 | - | 4qB | 22 | 4qB | 5 | 22.4±0.2 | 4qB | 23 |
|  | P02 | F | 18 | 12 | + | 379 | myopathic<br>change | ~18 | ~3 | 4qA | 3 | 4qA | 25 | - | - | - |
|  |  |  |  |  |  |  |  | ~98 | ~28 | 4qA | 28 | 4qA | 10 | - | - | - |
|  | P03 | F | 43 | 30 | + | 110 | mild<br>myopathic<br>change | ~18 | ~3 | 4qA | 3 | 4qA | 27 | - | - | - |
|  |  |  |  |  |  |  |  | ~38 | - | 4qA | 11 | 4qA | 34 | - | - | - |
|  | P04 | M | 27 | 11 | - | 1406 | myopathic<br>change | ~16 | ~3 | 4qA | 3 | 4qA | 18 | - | - | - |
|  |  |  |  |  |  |  |  | ~61 | ~17 | 4qB | 19 | 4qB | 42 | - | - | - |
|  |  |  |  |  |  |  |  | ~76 | ~21 | 4qA | 23 | 4qA | 30 | - | - | - |
|  | P05 | F | 41 | 31 | + | 95 | normal | ~12 | ~2 | 4qA | 2 | 4qA | 12 | - | - | - |
|  |  |  |  |  |  |  |  | ~38 | ~10 | 4qA | 11 | 4qA | 18 | - | - | - |
|  |  |  |  |  |  |  |  | >38 | - | 4qA | 17 | 4qA | 60 | - | - | - |
| Patient cohort 2 | P06 | F | 14 | 6 | - | 406 | myopathic<br>change | - | - | - | 2 | 4qA | 3 | 2.2±0.3 | 4qA | 32 |
|  |  |  |  |  |  |  |  | - | - | - | 15 | 4qB | 4 | 15.0±0.2 | 4qB | 36 |
|  |  |  |  |  |  |  |  | - | - | - | 27 | 4qA | 3 | 27.2±0.4 | 4qA | 5 |
|  | P07 | M | 23 | 18 | - | 871 | myopathic<br>change | - | - | - | 4 | 4qA | 5 | - | - | - |
|  |  |  |  |  |  |  |  | - | - | - | 20 | 4qA | 11 | - | - | - |
|  | P08 | M | 18 | 13 | - | 538 | myopathic<br>change | ~21.5 | ~5 | 4qA | 5 | 4qA | 6 | - | - | - |
|  |  |  |  |  |  |  |  | >63.5 | >17 | 4qA | 25 | 4qA | 9 | - | - | - |
|  | P09 | F | 33 | 19 | - | 269 | mild<br>myopathic<br>change | ~21.5 | ~5 | 4qA | 5 | 4qA | 33 | - | - | - |
|  |  |  |  |  |  |  |  | >63.5 | >17 | 4qA | 18 | 4qA | 26 | - | - | - |
|  | P10 | F | 39 | 28 | - | 176 | myopathic<br>change | ~15 | ~3 | 4qA | 3 | 4qA | 17 | - | - | - |
|  |  |  |  |  |  |  |  | >63.5 | >17 | 4qA | 28 | 4qA | 8 | - | - | - |
|  | P11 | M | 20 | 14 | + | 379 | myopathic<br>change | ~15 | ~3 | 4qA | 3 | 4qA | 16 | - | - | - |
|  |  |  |  |  |  |  |  | >63.5 | >17 | 4qA | 32 | 4qA | 12 | - | - | - |
|  | P12 | F | 15 | 10 | - | 514 | no data | ~12<br>~48.5 | ~2<br>15 | 4qA<br>4qA | 2<br>16 | 4qA<br>4qA | 10<br>18 | - | - | - |
|  | P13 | F | 53 | 32 | + | 341 | myopathic<br>change | ~18.5<br>>63.5 | ~4<br>>17 | 4qA<br>4qB | 4<br>18 | 4qA<br>4qB | 18<br>18 | - | - | - |
| Control | C01 | F | - | - | - | - | - | - | - | - | 19<br>47 | 4qB<br>4qA | 8<br>10 | 18.8±0.3<br>46.5±0.5 | 4qB<br>4qA | 22<br>13 |
|  | C02 | M | - | - | - | - | - | - | - | - | 18<br>20 | 4qA<br>4qA | 8<br>9 | 18.6±0.3<br>20.5±0.1 | 4qA<br>4qA | 7<br>8 |
|  | C03 | M | - | - | - | - | - | - | - | - | 13<br>22 | 4qB<br>4qB | 4<br>4 | 12.7±0.5<br>22.2±0.4 | 4qB<br>4qB | 20<br>22 |

**Figure 3.**
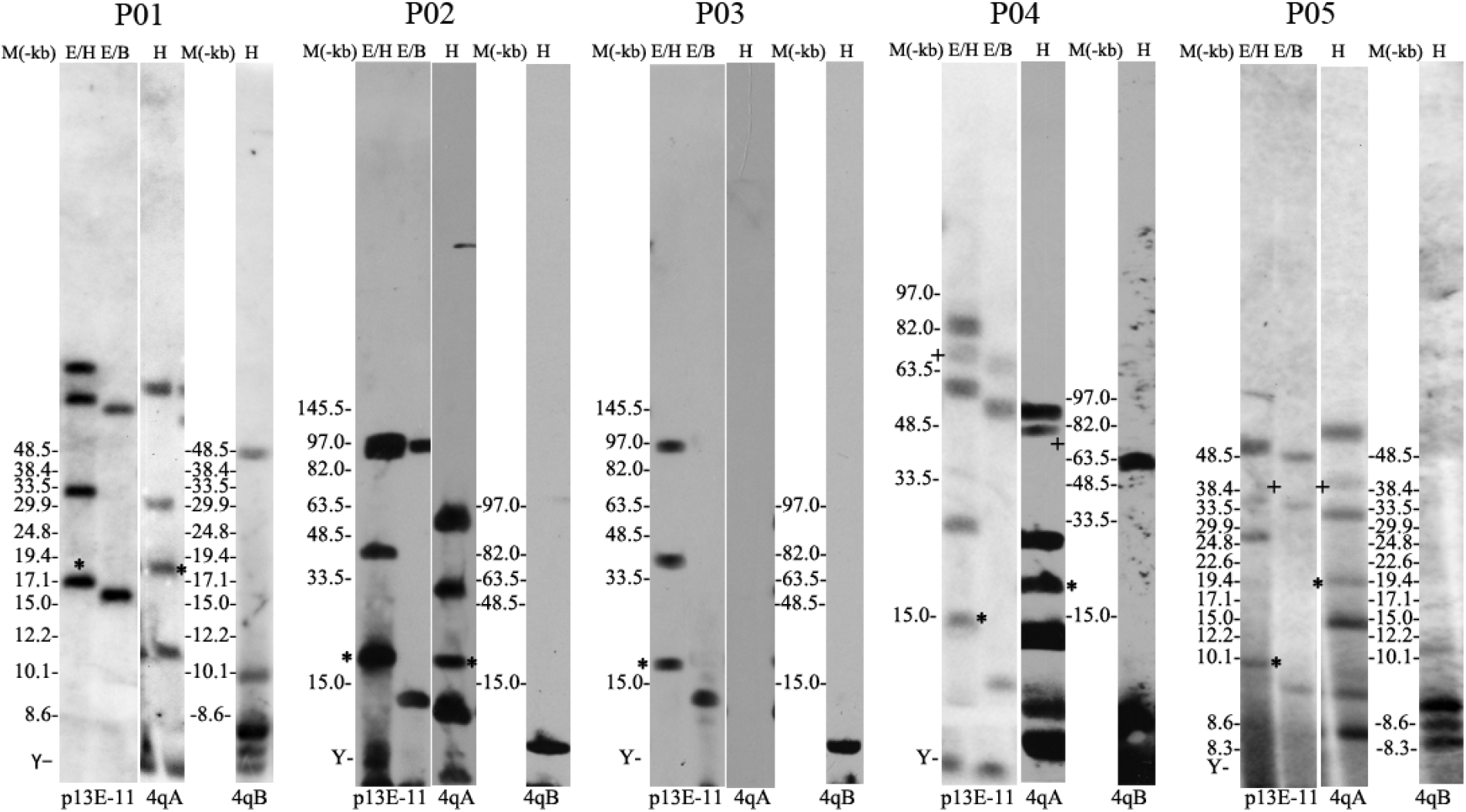
Molecular diagnosis of FSHD by Southern blot on samples from cohort 1. The results for cohort 2 is available in Supplementary Figure 2. E/H and p13E-11: Double digested with EcoRI/HindIII and then labeled with probe p13E-11, and all the 4q and 10q segments are illustrated. E/B and p13E-11: Double digested with EcoRI/BlnI and then labeled with probe p13E-11, and the 10q segments are digested so only 4q segments are illustrated. H and 4qA: digested with HindIII and then labeled with probe 4qA, and the 4qA alleles are illustrated. H and 4qB: digested with HindIII and then labeled with probe 4qB, and the 4qB alleles are illustrated. The star “*” denotes pathogenic allele with <10 repeat units and with a 4qA configuration. The plus sign “+” denotes somatic mosaic allele.

We next examined whether our methods can detect the presence of post-zygotic mosaicism on 4q35, since two patients were known to carry mosaicism from Southern blot. For example, for a patient (ID: P04), the presence of fluorescence labels can be directly visualized and counted in the Bionano Access browser, with 3, 19 and 23 repeats and the number of supporting reads as 18 (20%), 42 (46.7%) and 30 (33.3%), respectively. Since the three alleles were in 4qA, 4qB and 4qA configurations, respectively, we speculate that the allele with 3 repeats was due to a post-zygotic contraction of D4Z4 repeat units from the allele with 23 repeats, resulting in disease manifestation. We acknowledge that the read counts may be biased by the length of the reads and by the performance of the alignment algorithm. Furthermore, we also noticed that the repeat counts for the longer allele in P04 and P05 are slightly different (±1-2 repeats) from those inferred by Southern blot. Since Southern blot estimates repeat counts by band size in a gel, for alleles with larger number of repeats, it may be less quantitative than direct visualization of labels in optical mapping, and it cannot measure very long alleles (typically represented as more than a threshold such as “>38kb” or “>10 repeats” in diagnostic reports).

### Determining D4Z4 repeat copy number on 4q35 by Nt.BspQI enzyme labelling

Most published and ongoing human genetics studies use Nt.BspQI on the Bionano platform, partly due to its robustness and high sensitivity to recognize DNA motifs. Therefore, it would be ideal to develop methods that can utilize this enzyme as well, to determine the copy number of D4Z4 repeat units. Our *in silico* digestion analysis on the reference genome GRCh38 demonstrated that it is possible to differentiate reads originating from 4q35 and 10q26 (Figure 2B). Although the reference genome GRCh38 contains two Nt.BspQI labels within the D4Z4 repeat region, we do not observe them in real data, possibly due to the inclusion of a very rare allele or due to genome assembly errors in GRCh38. Instead, based on recognition sites surrounding the D4Z4 repeat array, we can infer the length of the D4Z4 array purely based on the distance between flanking regions with fluorescence labels (Figure 2B). Note that even if a very rare allele does include enzyme recognition site within the D4Z4 repeat, it will not negatively impact our calculation, as our method estimate the length of the repeat based on fluorescence label pattern of the flanking regions. Additionally, although the human reference genome GRCh38 only contains sequences for the 4qB configuration, our empirical analysis and previous report [23] demonstrated that the 4qA/4qB allele type of the repeat units can be confidently assigned by the presence of a 5-label or 3-label array distal to the D4Z4 repeat array. We additionally note that the 4qA sequence was recently added into CRCh38 patch 7 (KQ983257.1), as illustrated in Figure 1 and Figure 2.

To further examine this possibility, we performed optical mapping on selected patients using the Nt.BspQI enzyme. For example, for the patient (ID: P01) in **Supplementary Figure 1**, the distance between two flanking segments of the D4Z4 clusters can be used to quantify the number of repeats. Unlike our analysis using the Nb.BssSI enzyme where the enzyme recognition site is directly located within the D4Z4 repeat unit, we can assign a quantitative (floating-point) repeat count to each DNA molecule that spans the region of dissimilarity, since the enzyme recognition sites are outside of the repeat regions. The information from all reads can be compiled together to reach an estimate of the repeat counts for both alleles (4 and 22 repeats, respectively), and our results were completely consistent with the results obtained from the Nb.BssSI enzyme.

### Additional validation on a second patient cohort and control subjects

To further validate the method in diagnostic testing settings, we analyzed a second cohort of 8 individuals suspected to have FSHD, from two separate institutions. These patients all had typical clinical manifestations consistent with FSHD (Table 1). However, they were not previously subject to any diagnostic testing of FSHD. We performed the genetic analysis on the Bionano Saphyr single-molecule optical mapping platform. We obtained positive results on all the patients, and found that one patient (ID: P06) had post-zygotic mosaicism with 2, 15 and 27 repeats (Figure 4). We validated this result using both the Nb.BssSI enzyme and Nt.BspQI enzyme. Since the allele with 15 and 27 repeats had 4qB and 4qA configuration, respectively, we can further infer that the post-zygotic contraction of D4Z4 repeats occurred on the allele with 27 repeats in 4qA configuration, which represented a dramatic decrease of D4Z4 copy number on this allele. Finally, we obtained Southern blot based diagnosis from an independent diagnostic lab on 6 individuals for whom sufficient DNA is available (**Supplementary Figure 2**). The estimated repeats from Southern blot are highly consistent with those inferred from optical mapping (Table 1, **Supplementary Table 2**).

**Figure 4.**
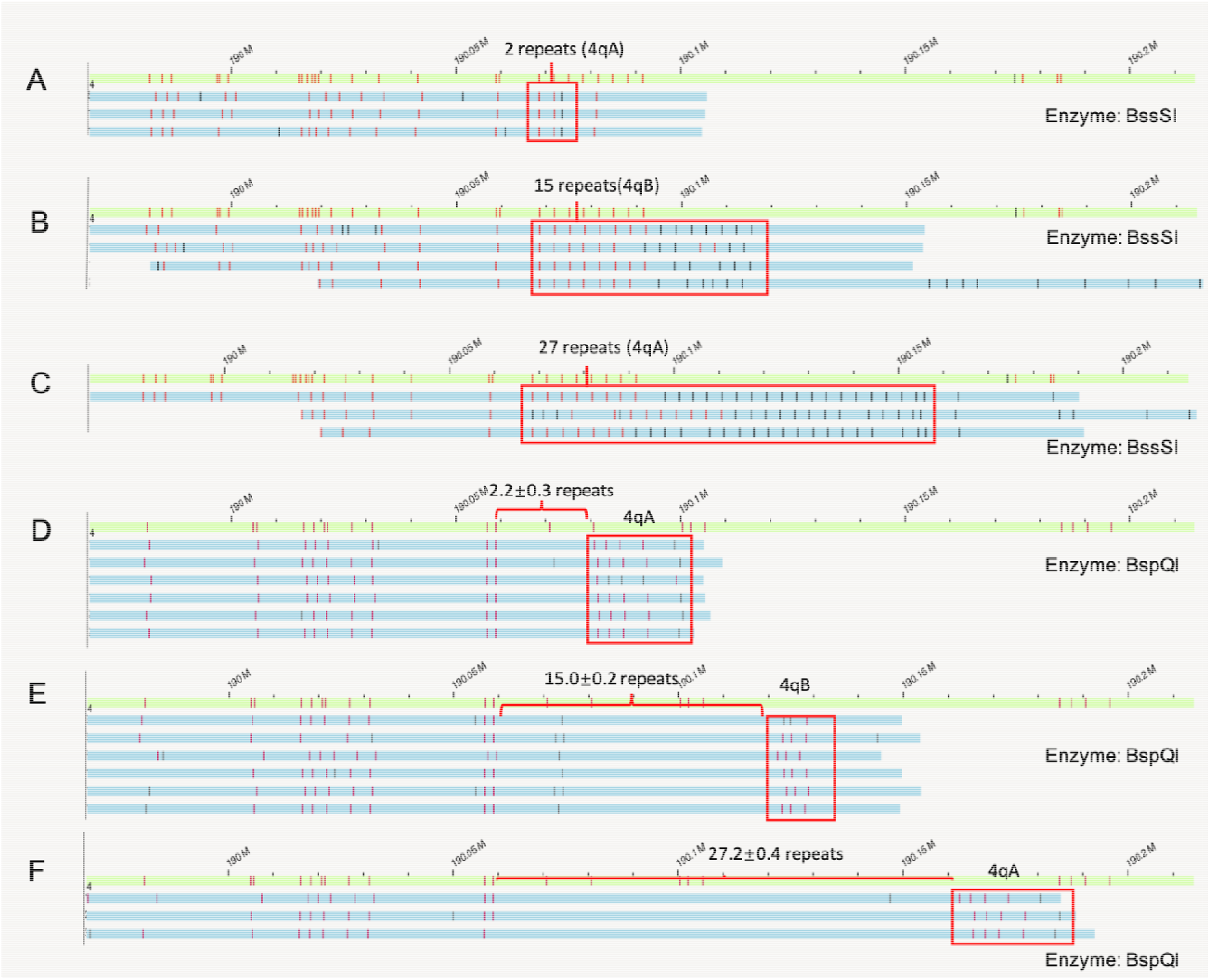
Determination of somatic mosaicism by Nb.BssSI (A/B/C) and Nt.BspQI (D/E/F) enzymes on a patient with 2, 15 and 27 repeats (ID: P06). For both enzymes, our computational pipeline accurately identified the presence of somatic contraction, and determined that the contraction occurs on the parental allele carrying the 4qA configuration. For repeat quantification using the Nt.BspQI enzyme, mean ± SD (standard deviation) is annotated in the figure. Vertical bars represent labels of enzyme recognition sites.

In addition to patient cohorts, we evaluated our approach on several control subjects without FSHD and without a family history of FSHD. We also downloaded publicly available Bionano genome mapping data sets on the NA12878 subject (ID: C01). The results from both the Nb.BssSI enzyme and Nt.BspQI enzyme were consistent with each other for all the subjects (Table 1), suggesting that the method can work on healthy human populations with larger number of D4Z4 repeat units.

Finally, we evaluated whether our method can be applied to an earlier generation of the single-molecule optical mapping platform, the Bionano Irys platform. We performed additional analysis on three control families from the 1000 Genomes Project (Table 2). With one exception, we found that the D4Z4 repeat unit numbers of the offspring and the parents in all three families were consistent with Mendelian inheritance, demonstrating high accuracy of the method. Additionally, the results from the Nb.BssSI enzyme and Nt.BspQI enzyme were also consistent with each other, except for three subjects who carried large unit numbers. However, we also note that for three subjects, only one allele was quantified using the data sets of Nb.BssSI enzyme, but both alleles can be quantified using the data sets of Nt.BspQI enzyme. Additionally, the 4qA/4qB configuration for one subject is different between the results generated by the Nt.BspQI and Nb.BssSI enzyme. The largest number of units detected in our study is 49. Thus, we caution that optical mapping on the older Irys platform may be less effective for human subjects carrying >50 D4Z4 repeat units, given the size limitations of typical DNA extraction and optical mapping experiments on the Irys platform. In summary, we caution that the Irys platform may limit the performance, due to generally lower throughput (lower coverage) and shorter DNA fragments during DNA preparation, and may generate incorrect 4qA/4qB allelic configurations.

**Table 2.** Analysis of Bionano Irys genome mapping data sets of three families.

|  | ID | Ethnicity | Relationship | Irys optical mapping<br>(Nb.BssSI enzyme) |  |  | Irys optical mapping (Nt.BspQI<br>enzyme) |  |  |
| --- | --- | --- | --- | --- | --- | --- | --- | --- | --- |
|  |  |  |  | Units | Allele | Read<br>Count | Units | Allele | Read<br>Count |
| Family 1 | HG00512 | SOUTHERN<br>HAN<br>CHINESE | father | 19 | 4qA | 3 | 19.8±0.5 | 4qB | 7 |
|  |  |  |  | - | - |  | 40.4±0.3 | 4qA | 3 |
|  | HG00513 | SOUTHERN<br>HAN<br>CHINESE | mother | 11 | 4qA | 6 | 10.4±0.2 | 4qA | 8 |
|  |  |  |  | - | - |  | 33.1±0.3 | 4qA | 11 |
|  | HG00514 | SOUTHERN<br>HAN<br>CHINESE | daughter | 11 | 4qA | 6 | 10.6±0.2 | 4qA | 9 |
|  |  |  |  | 19 | 4qA | 5 | 19.6±0.3 | 4qA | 10 |
| Family 2 | HG00731 | PUERTO<br>RICAN | father | 20 | 4qB | 5 | 19.9±0.5 | 4qB | 7 |
|  |  |  |  | 40 | 4qA | 3 | 40.5±0.3 | 4qA | 3 |
|  | HG00732 | PUERTO<br>RICAN | mother | 17 | 4qB | 6 | 17.2±0.1 | 4qB | 33 |
|  |  |  |  | 32 | 4qB | 4 | 32.1±0.3 | 4qB | 18 |
|  | HG00733 | PUERTO<br>RICAN | daughter | 17 | 4qB | 7 | 17.3±0.3 | 4qB | 12 |
|  |  |  |  | 40 | 4qA | 2 | 40.1±0.4 | 4qA | 7 |
| Family 3 | GM19238 | YORUBA | mother | 25 | 4qA | 2 | 25.6±0.3 | 4qA | 10 |
|  |  |  |  | - | - | - | 49.1±0.7 | 4qA | 13 |
|  | GM19239 | YORUBA | father | 9 | 4qB | 5 | 9.2±0.1 | 4qB | 14 |
|  |  |  |  | 36 | 4qA | 2 | 36.6±0.2 | 4qA | 13 |
|  | GM19240 | YORUBA | daughter | 25 | 4qA | 3 | 25.9±0.3 | 4qA | 5 |
|  |  |  |  | 36 | 4qA | 4 | 36.2±0.4 | 4qA | 7 |

## DISCUSSION

In this study, we evaluated the technical feasibility of using NanoChannel-based optical mapping to characterize D4Z4 repeat numbers and allelic configurations in FSHD. We demonstrated that this method can accurately quantify the number of repeats, can differentiate the DNA fragments from 4q35 and 10q26, and can quantify the mosaic levels of repeats when one allele has a post-zygotic contraction of D4Z4 repeat units. We concluded that optical mapping is a viable approach for quantifying D4Z4 repeats in FSHD and may be applied in clinical diagnostic settings once more validations are performed in the future.

Compared to conventional optical mapping approaches, the Nanochannel-based optical mapping has several clear advantages. First, by stretching the DNA molecules as linear molecules and going through massively parallel Nanochannels, the resolution and throughput are much higher than conventional optical mapping approaches that spread labeled DNA molecules on glass slides in semi-controlled fashion. Additionally, when Nb.BssSI enzyme is used, the labels (one for each D4Z4 unit) can be directly visualized in optical mapping platform. As illustrated in a previous study using optical mapping on FSHD [24], the ability to visually check and count repeats is a major advantage over Southern blot and FISH combing. A second advantage is the flexibility to switch to different enzymes to allow detection of different patterns. In our study, we demonstrated that Nb.BssSI enzyme is a preferred choice for FSHD since an enzyme recognition site is directly located within the D4Z4 repeat unit, but we also developed methods to quantify the repeat number with the Nt.BspQI enzyme labeling. Recently, molecular combing is also used to detect the copy number of D4Z4 repeat units [16]. However, molecular combing only detects the approximate length of the D4Z4 array and the repeat number is inferred from the length. In comparison, we can calculate repeat numbers directly using the Nb.BssSI enzyme, while also estimate the length of D4Z4 array using the Nt.BspQI enzyme in a similar fashion as molecular combing. We also recognize that long-read sequencing technologies, such as PacBio SMRT (single-molecule real-time) sequencing and Oxford Nanopore sequencing, may also be used in the molecular diagnosis of FSHD. However, these sequencing techniques usually produce data with read length N50 < 20kb. Ultra-long Nanopore sequencing with a special library construction procedure may produce read length N50 of 100 kb or higher, but it requires much more input DNA (~10 μg) with lower data yield. Another advantage of the Saphyr platform is that in addition to quantifying the copy number of D4Z4 repeat units, it will also enable the *de novo* assembly of a human genome and the genome-wide identification structural variants [25]. Therefore, optical mapping can serve a dual purpose of identifying structural variants that may be relevant to the phenotypic presentations in the patient, especially when FSHD diagnostic testing yields negative results.

We also recognize that there are several disadvantages of optical mapping, in comparison to conventional approaches such as Southern blot, in the contexts of molecular diagnosis of FSHD. First, although Southern blot may not be very accurate (especially for alleles with more than 17 repeats), for the purpose of molecular diagnosis, it can roughly tell the number and enable the diagnosis of FSHD when an allele with 1-10 repeat is observed. This straight-forwardness may be one of the reasons for Southern blot to be commonly used for the diagnosis. Second, optical mapping is not yet cost-effective in comparison: there is an initial capital cost of a few hundred thousand dollars to establish the platform itself (including the computing hardware), and each subsequent flow cell currently costs ~$500 even when purchased in batch. We note that it is possible to lower the cost in future versions of the flow cell by assaying >10 samples together (currently, it is limited to two samples), and that optical mapping can perform genome-wide survey of structural variants that may be relevant to disease diagnosis. Furthermore, although the Saphyr platform can run two human genomes in one day in an automated fashion (approximately ~300Gb of data for each genome), the sample preparation itself takes approximately 1 day with extensive manual labor (see Methods), and the data analysis and visualization can take half a day. Southern blot based diagnosis typically takes one week, but it has the clear advantage that more than 10 samples can be analyzed on the same gel (and it is possible to run 4 gels in one week), allowing the simultaneous analysis of a large number of samples. Third, the number of reads encountered in optical mapping is highly size dependent, while this is not an issue using high quality DNA (from agarose plugs) in a Southern blot; therefore, optical mapping may be less accurate when estimating the proportion of mosaicism when several alleles differ substantially in repeat counts.

There are several technical limitations of optical mapping that we wish to discuss here. First, our study focused on the molecular characterization of patients with FSHD, and only included a very small number of unaffected control subjects. Currently the average length of reads from the Saphyr platform is about 350kb, even though the DNA molecules that were assayed in our study can range from 100kb to over 1Mb. Therefore, very long non-pathogenic D4Z4 repeats could not be accurately quantified by optical mapping. Indeed, if an allele has 50 units in control subjects, then the length of the repeat region would be 50*3.3kb=165kb; given that typical average size of an optical mapping is ~350kb, many of the reads from optical mapping may not be able to cover the “region of dissimilarity”, so the effective coverage at the D4Z4 region will be much less than the genome-wide coverage (which is usually ~100X per flowcell). We illustrate the relationship between coverage and read length in **Supplementary Table 3**: it is clear that samples with similar whole-genome coverage can vary greatly when focusing on longer reads: some samples such as P01 and P08 has very low effective coverage for reads>300kb in comparison to other samples. This is not a problem for diagnosing FSHD *per se*, since <10 repeats are pathogenic, but it may pose a problem for population-scale analysis of D4Z4 repeats since it gives an upper bound of the number of repeats that can be detected by the platform. Second, for long-read platforms, there will be allelic biases where longer alleles tend to be present with lower number of reads than shorter alleles. Given the size distribution of all reads genome-wide calculated by the Bionano software, it is possible to estimate coverage biases of the two alleles. To further examine this issue, we have also compared the number of reads supporting shorter versus longer D4Z4 alleles for all subjects (**Supplementary Table 2**). Except the 3 individuals with post-zygotic mosaicism, among the 14 subjects, 5 have less reads covering the shorter alleles than the longer alleles; therefore, this potential bias does not appear to be a main concern. One reason why the length bias is not high may be due to the need to find very long flanking alignment around the D4Z4 repeats as shown in Figure 2, so that the total length of aligned regions between the two alleles are comparable. Third, if there is zero repeat in an allele, then it will be hard to tell heterozygosity from homozygosity; in our samples we did not observe such events to evaluate this possibility, but they are likely to occur in the population. Fourth, the optical mapping method cannot distinguish 4qA-L from 4qA/4qB. To the best of our knowledge, there is little clinical implication whether 4qAL or 4qA is present, since they both show phenotypes of FSHD when repeat number is 1-10. We note that the GRCh38.p7 release also includes KQ983258.1 as patch scaffold providing representation for the variant 4qA-L haplotype which is slightly (~1.6kb) longer than the 4qA haplotype.

We also wish to discuss several limitations of the current study design. First, with the exception of patient P02 and P03 (offspring and mother), we were not able to obtain parental data for patients under the study. As previously reviewed [26], several studies demonstrated that *de novo* repeat contraction may account for a surprisingly high percentage of FSHD patients (10%-33%) [27, 28], and this high incidence can be partly explained by the presence of parental mosaicism for 4q short alleles that has been reported in 19% of *de novo* cases [29, 30]. Lemmers et al [31] demonstrated that somatic mosaicism in FSHD patients goes largely undetected using the standard diagnostic technique, indicating that linear electrophoresis is unsuitable to identify mosaic patients, yet pulsed-field gel electrophoresis (PFGE)–based method can accurately reveal somatic mosaicism in patients and parents. Among the three patients with post-zygotic mosaicism in our study (P04, P05 and P06), one of them (P05) had a family history of FSHD and a relatively late onset at age 31. Detailed examination of medical records showed that P05 was a female patient who was referred to the clinic due to mild symptoms and due to a confirmed diagnosis of an offspring with early-onset FSHD. Note that we were unable to determine the genetic origin of the pathogenic mutation in P05 due to the lack of parental data; similarly, we were unable to determine whether the contracted repeat number differs between P05 and her offspring as the offspring did not consent in this study. Nevertheless, this case represented an interesting example where post-zygotic mosaicism was inferred in a patient suspected to carry germline mosaicism, corroborating previous reports that a substantial fraction of mosaic parents with germline mosaicism in oogenesis may have been overlooked [30]. One additional limitation of the current study is that we used blood samples for molecular diagnosis, rather than muscle biopsy from affected areas. However, in practice, it is generally not feasible and not desirable to take a muscle biopsy for DNA analysis, therefore limiting our ability to compare results between blood and tissues in the current study. Expanding muscle derived cells *in vitro*, such as satellite cells or iPS cells, might be helpful in determining the degree of mosaicism in affected muscles.

In conclusion, we established the technical feasibility of using Bionano Genomics’s Saphyr platform to perform molecular diagnosis of FSHD, and discussed a number of advantages, limitations and possible modifications that may improve the detection accuracy and reliability. With the ever decreasing cost of performing genome mapping on the single-molecule optical mapping platform, and the recent introduction of Direct Label and Stain technology, we expect that this method may be widely applied in research and clinical settings of FSHD, and may potentially expedite the genetic studies on this devastating disease. Lastly, this study may serve as a model, which demonstrated how the workflow can be applied to other rare diseases involving complex genomic structural changes.

## MATERIALS AND METHODS

### Sample selection

The study was approved by the Institutional Review Board of the Peking Union Medical College of the Chinese Academy of Medical Sciences (IRB #JS-1316). Our primary patient cohort consists of five patients, including two from the same family (P02 and P03). Four of the patients have a family history of FSHD, but one patient (P04) is sporadic (Table 1). All the patients had a molecular diagnosis of FSHD by Southern blots, and two of the patients were known to carry mosaicism based on Southern blots, though the exact fraction and the parental origin of the chromosome with post-zygotic contraction cannot be determined. We stress here that the Southern blot and optical mapping on this cohort were blindly performed in two separate institutions without information from each other, yet the final results are consistent. Our second patient cohort consists of eight patients suspected to have FSHD, but without a prior molecular diagnosis. We performed Southern blot based diagnosis on six patients with available DNA samples, as a validation assay after we obtained results from optical mapping. Additionally, existing data on two unaffected adult subjects without FSHD, as well as the publicly available data one individual and three families, were included in the study as negative controls.

### Southern blot and PFGE-based DNA analysis

We followed previously published protocols for Southern blot [10, 32]. All diagnostic testing by Southern blots were performed at the Department of Neurology, First Affiliated Hospital of the Fujian Medical University. For each sample, one portion of DNA samples was double digested with EcoRI/HindIII and with EcoRI/BlnI, yet another portion of DNA samples was digested only with HindIII. Then the digested DNA was separated by pulsed field gel electrophoresis (PFGE). After electrophoresis, the DNA was transferred to a Nytran XL membrane and hybridized with the probes p13E-11, 4qA, 4qB, respectively. Finally, the blots were exposed to obtain images for further manual analysis of repeat numbers.

### High molecular weight (HMW) DNA isolation for optical mapping

Fresh blood samples were collected in EDTA stabilized anti-coagulative tube, adequately mixed and stored at 4°C promptly. Samples that cannot be processed within 5 days after collection were stored at - 80°C. HMW DNA was extracted from either 1 ml frozen or fresh blood, following manufacturer’s guidelines (Bionano Prep Blood DNA Isolation Protocol, Bionano Genomics, #30033) with slight modifications. Briefly, red blood cells were lysed by RBC lysis solution (Qiagen) and white blood cells (WBC) left were pelleted. After centrifugation, WBC were re-suspended in cell buffer (Bionano Genomics, USA), and then the WBC solutions were embedded into 2% agarose plugs (CHEF Genomic DNA Plug Kit, Bio-Rad) to avoid fragmentation of long DNA molecules, with overnight lysis at 50°C in lysis buffer (Bionano Genomics, USA) with Puregene Proteinase K (Qiagen). The volume ratio (v/v) of WBC and agarose were typically 1.5:1, 2:1, 4:1, 8:1 and 16:1, in order to obtain DNA with proper concentration. (We typically do a quick DNA extraction before embedding the cells into the plug, and we choose a ratio that yields ~2.2ug DNA per plug which corresponds to approximately 6×10^5^ cells.) The agarose plugs were washed with Tris-EDTA buffer the following day and digested at 43°C with GELase™ Agarose Gel-Digesting Prep (2 unit/µl, Thermo Fisher) for 50 min. Extracted HMW DNA was purified via drop dialysis using Millipore membrane filters (EMD Millipore, USA) placed on Tris-EDTA buffer for 3 hours.

DNA quantification was carried out using Qubit dsDNA assay BR kits with a Qubit 2.0 Fluorometer (ThermoFisher Scientific). The integrity of HMW DNA was detected by pulsed-field gel electrophoresis (Pippin Pulse, Sage Science). Only the DNA samples with concentration between 30-100ng/ul and sufficient molecular mass were used in the following DNA labeling experiment.

### DNA labeling and chip loading

The DNA labeling experiment (also referred to as “NLRS”) consists of four sequential steps (Nick, Label, Repair and Stain), and was performed strictly following manufacturer’s guidelines (Bionano Prep™ Labeling - NLRS Protocol, Bionano Genomics, #30024). In short, 300 ng of purified HMW DNA was nicked by nicking endonucleases Nb.BssSI (New England BioLabs) in 10X Buffer 3.1 (Bionano Genomics) at 37 °C for 2 hours. Using Taq polymerase (New England BioLabs), the nicked DNA was labeled at 72 °C for 1 hour by fluorophore-labeled nucleotides mixed in 10X Labeling mix (Bionano Genomics). In the third step, labeled DNA was repaired with Taq ligase (NEB) at 37 °C for 30 min to restore integrated double strands DNA. In the last step, the DNA backbone was stained overnight in a dark environment at 4 °C for visualization and size identification. DNA quantification was carried out using Qubit dsDNA assay HS kits with a Qubit 2.0 Fluorometer (ThermoFisher Scientific) and only DNA samples with concentration between 4-10ng/ul were chosen to be loaded in the next step.

Labeled DNA was loaded on the Saphyr chip and pushed by low-voltage electric field into pillar region and Nanochannel in the Bionano Saphyr instrument. After the mapping procedure begins, the fluorescently labeled DNA molecules were imaged by the Saphyr instrument. Generally, after 24-36 hour, each sample can generate 320-480 Gbp data for each flowcell, which was used for further data analysis.

### Downloading data sets from publicly available resources

The data sets on the NA12878 genome were downloaded from Bionano Genomics’s website (https://bionanogenomics.com/library/datasets/). The data sets of the three family trios in the 1000 Genomes Project were also provided by Bionano Genomics.

### Bioinformatics approaches to analyze optical mapping data

The detailed description of the bioinformatics algorithms is provided below. In particular, we addressed the challenges in the differentiation of 4qA with 4qB alleles, the differentiation of 4q35 and 10q26 segmental duplications, the quantification of repeat numbers with different enzymes that may or may not have recognition sites within D4Z4 repeats.

### Data pre-processing and alignment

The raw output files from the Bionano Saphyr mapping platform and Irys platform were in BNX formats. Each file contained molecule and label information and quality scores per channel identified during a Bionano run. Where necessary, for each subject, we combined several BNX files into one BNX file. We next performed a basic filtering of the BNX file using all default parameters suitable for human genome, including the 150kb length cutoff and the label SNR filter. To assess the quality of each of the Bionano runs, we performed Molecular Quality Report using default parameters and examined the results by comparing to manufacturer recommended values. Additional details can be found at https://bionanogenomics.com/wp-content/uploads/2017/05/30175-Rev-A-Bionano-Molecule-Quality-Report-Guidelines.pdf.

For each enzyme, we performed *in silico* digestion of the human reference genome (GRCh38) to generate the reference map for that particular enzyme (that is, the reference CMAP file). We then mapped the BNX files to the reference CMAP file, using the Bionano-Solve (version 3.1), accessed from https://bionanogenomics.com/support/software-downloads/. Slight modifications were made to the source code align_bnx_to_cmap.py to change all the hard-coded path name in the software. The results include three file types: xmap, r.cmap and q.cmap. They represent the alignment file, the reference label file and the query label file, respectively. A custom script was developed to extract specific regions of alignments from whole-genome mapping to speed up subsequent data analysis and visualization.

### Determination of repeat copy number and 4qA/4qB configuration by Nb.BssSI enzyme

We developed a simple yet effective approach for quantifying repeat units and determining 4qA/4qB configurations using Nb.BssSI-based labels. Through the analysis of an empirical mapping data set on human samples, which was already normalized through a scaling factor to account for the variation of DNA migration rates during Bionano data acquisition, we calculated that the length of D4Z4 repeat unit (hereafter referred to as *r*) is 3299.4bp ± 144.00bp (mean ± standard deviation), the distance of 4qA to the D4Z4 repeat unit (hereafter referred to as *q*) is 1758.79bp ±61.15bp and the distance to the first label immediately after 4qA to the 4qA label (hereafter referred to as *s*) is 7889.79bp ± 240.93bp. These observations suggest that the three measures (*r*, *q*, *s*) are quite distinct from each other and their variances are small enough so that they can be easily differentiated in real data. Based on these empirical observations, we set a coefficient of variation (hereafter referred to as *t*) upper bound of 0.05, which is a measure of relative variability as the ratio of the standard deviation to the mean (the empirical observations had *t* values of 0.044, 0.035 and 0.031 for the three measures, respectively).

Our algorithm accounts for the relatively high error rates in the label data. Based on technical documentation from Bionano (https://bionanogenomics.com/wp-content/uploads/2017/05/30175-Rev-A-Bionano-Molecule-Quality-Report-Guidelines.pdf), the percentage of unaligned labels in molecules relative to number of labels in molecules and the percentage of unaligned reference labels relative to number of reference labels are generally less than 15% and 21%, respectively, for a mapping data set with reasonable quality. Our algorithm first examines reads that are mapped to the 4q35 target region (Figure 2) with sufficient number of labels in the “region of dissimilarity” to ensure that the reads originate from 4q35 rather than 10q26. For each read, we denote the position of the first label within D4Z4 region as *l*_*1*_ for simplicity, and all following labels as *l*_*2*_, *l*_*3*_ up to *l*_*n*_ (the last label in the read). We then calculate a distance vector between all adjacent labels as *d* = [*d*_*1*_, *d*_*2*_, *d*_*3*_,…, *d*_*n-1*_] = [*l*_*2*_ − *l*_*1*_, *l*_*3*_ − *l*_*2*_, *l*_*4*_ − *l*_*3*_,…, *l*_*n*_ − *l*_*n-1*_]. Based on each *d*_*i*_ (1≤ *i* ≤ *n-1*) in the vector, we classify each label into one of five categories of events: (1) one additional D4Z4 repeat, when *r*(*1* - *t*) ≤ *d*_*i*_ ≤ *r*(1 + *t*); (2) one or two false negative labels (that is, expected label is missing from reads), when *r*(1 + *t*) < *d*_*i*_ ≤ 3*r*(1 + *t*); (3) one or two false positive labels (that is, extra label is present in reads), when *r*(1 - *t*) < *d*_*i*_ +*d*_*i+1*_ ≤ *r*(1 + *t*) for one extra label, or when *r*(1-*t*) < *d*_*i*_+*d*_*i+1*_+*d*_*i+2*_ ≤ *r*(1 + *t*) for two extra labels; (4) 4qA is encountered, when *q*(1 - *t*) ≤ *d*_*i*_ ≤ *q*(1 + *t*) and when *s*(1 - *t*) ≤ *d*_*i*+1_ ≤ *s*(1 + *t*); (5) a label outside of D4Z4 repeat region is encountered, or an exception is encountered when the first four criteria are not met. Optionally, the prior probability for the five events can be determined from genome-wide estimate of false positive and false negative rates when mapping all reads to the reference genome GRCh38, to assign reads into the five categories based on posterior probability. When more than one consecutive category 5 events are encountered, the label counting for this read ends and the number of estimated repeat counts is recorded. For each sample, the results for all reads are then tallied in a histogram. A peak-calling algorithm is used to classify one peak (homozygous, which is rare), two peaks (heterozygous) or three peaks (post-zygotic mosaicism), and quantify the number of repeat units corresponding to each peak. We also note that although we have not encountered this situation in practice, it is possible that all reads spanning this region have only category 5 events, when the patient carries zero copies of D4Z4 repeat units in both alleles.

### Determination of repeat copy number and 4qA/4qB configuration by Nt.BspQI enzyme

Although less straightforward, we also developed a simple yet effective approach for quantifying repeat units and determining 4qA/4qB configurations using Nt.BspQI-based labels. Based on the published sequence of D4Z4 (GenBank: D38024.1) [33], we determined the position of D4Z4 repeat units on 4q35 in the human reference genome GRCh38 as shown in **Supplementary Table 4** (we note that a previous study [23] incorrectly determined repeat units in GRCh38). However, with one possible exception, analysis of a large number of real data sets failed to detect Nt.BspQI labels in repeat unit #2 and #5, suggesting that GRCh38 may have included a very rare allele or may contain assembly errors in this region. Similarly, we determined the position of D4Z4 repeat units on 10q26 (**Supplementary Table 5**) and on KQ983257.1 (**Supplementary Table 6**), and illustrated the presence of a genome assembly gap on 10q26 that incorrectly created two separate repeat arrays in GRCh38 (GRCh37 is not influenced by this issue as shown in Figure 2).

Based on *in silico* enzyme digestion of GRCh38 by the Nt.BspQI enzyme, we found that there is a restriction enzyme recognition site 7,271bp upstream of the first D4Z4 repeat unit in GRCh38 (CMAP coordinate as chr4: 190058870), and that there is another restriction enzyme recognition site 7,860bp downstream of the last D4Z4 repeat unit in GRCh38 (CMAP coordinate as chr4: 190100364). Therefore, when we can anchor a read to the two enzyme recognition sites, we can calculate the distance between two labels as *x*, and estimate the number of D4Z4 repeat units as *y*=(*x*-7271-7860)/3300. Unlike our analysis on Nb.BssSI, the estimate *y* here is a floating point value rather than an integer. When multiple reads are mapped to the same region, we can calculate the value of *y* to yield a best estimate of the number of repeat units. As previously discussed [23], based on empirical evidence, the 4qA and 4qB configuration can be differentiated by the presence of five labels (4qA) or three labels (4qB) downstream of D4Z4 repeat region.

### Visualization and manual examination of results

We used the Bionano Access software for visualization of genome mapping and manual examination of results. The software was obtained from https://bionanogenomics.com/support-page/bionano-access/. It is a node.js web application that can communicate with a remote server, but can also run in standalone mode to perform visualization of results.

We extracted a subset of reads that mapped to 4q35 and 10q26 regions and performed manual examination of their alignment. The results are generally consistent with computational results, further suggesting that the method is highly reliable.

## DECLARATIONS

### Ethics approval and consent to participate

The study was approved by the Institutional Review Board of the Peking Union Medical College of the Chinese Academy of Medical Sciences. All individual participants involved in this study have consent to participate.

### Consent for publication

All individual participants have consent for publication.

### Competing interests

P.L., F.L., F.Y., J.Z., and D.W. are employees and K.W. was previously a consultant of Grandomics Biosciences.

### Author contributions

YD, LC and KW planned the study. YD, FY, ZW provided clinical diagnosis of the FSHD patients and performed sample selection, preparation, southern blotting and optical mapping. PL, FL, LF developed the computational pipeline and analyzed the single-molecule optical mapping data. YH, SH, JZ, DW advised on the execution of the study and data interpretation. LC and KW are responsible for the overall content and are corresponding authors.

### Funding

This study is in part supported by the CAMS Innovation Fund for Medical Sciences (CIFMS) project number 2016-I2M-1-002.

## Acknowledgements

We thank the patients with FSHD who participated in this study to develop novel diagnostic tests using genomic technologies. We also thank the genetic counselors and clinical geneticists who interviewed the patients and collected samples. We thank technical team at Bionano Genomics to generate control data sets from the 1000 Genomes Project, provide technical support and offer suggestions. This study was partially supported by institutional resources at the Peking Union Medical College of the Chinese Academy of Medical Sciences.

