## Supplementary Materials for "Single-molecule optical mapping enables quantitative measurement of D4Z4 repeats in facioscapulohumeral muscular dystrophy (FSHD)"

### Supplementary Table

**Supplementary Table 1. Additional descriptions on the clinical manifestation of several patients with FSHD in the study.**

| ID | Clinical Manifestation |
|---|---|
| P01 | facial muscle weakness(orbicularis oris), mild upper arms weakness, Beevor(+) |
| P02 | facial muscle weakness(orbicularis oculi and orbicularis oris), upper arms weakness, scapular winging, Foot drop, Beevor(+) |
| P03 | facial muscle weakness(orbicularis oculi and orbicularis oris), mild upper arms weakness, scapular winging, Beevor(+) |
| P04 | facial muscle weakness(orbicularis oculi and orbicularis oris), upper arms weakness, scapular winging, Foot drop, Beevor(-) |
| P05 | facial muscle weakness(orbicularis oculi and orbicularis oris), mild scapular winging, Beevor(-) |
| P06 | facial muscle weakness(orbicularis oculi and orbicularis oris), upper arms weakness, scapular winging, Foot drop, Beevor(+) |
| P07 | facial muscle weakness(orbicularis oculi and orbicularis oris), upper arms weakness, scapular winging, Beevor(+) |
| P08 | facial muscle weakness(orbicularis oculi and orbicularis oris), upper arms weakness（asymmetric）, scapular winging,foot drop, Beevor(-) |
| P09 | facial muscle weakness(orbicularis oculi and orbicularis oris), upper arms weakness（asymmetric）, scapular winging,foot drop, Beevor(+) |
| P10 | facial muscle weakness(orbicularis oculi and orbicularis oris), upper arms weakness, scapular winging,foot drop, Beevor(+) |
| P11 | facial muscle weakness(orbicularis oris), upper arms weakness（asymmetric）, scapular winging,foot drop, Beevor(+) |
| P12 | facial muscle weakness(orbicularis oculi and orbicularis oris), upper arms weakness（asymmetric）, scapular winging, Beevor(+) |
| P13 | facial muscle weakness(orbicularis oris), upper arms weakness（asymmetric）, scapular winging, Beevor(+) |

**Supplementary Table 2. A list of patients and control subjects assayed by Bionano Saphyr platform in the current study.** An abbreviated version of this table can be found in Table 1. EMG: Electromyography, CK: creatine kinase.

| | ID | Sex | Age (years) | Onset (years) | Family history | CK (U/L) | EMG | Southern Blot (4q35) | | | Southern Blot (10q26) | | | Optical mapping (Nb.BssSI enzyme, 4q35) | | | Optical mapping (Nb.BssSI enzyme, 10q26) | | | Optical mapping (Nt.BspQI enzyme, 4q35) | | | Optical mapping (Nt.BspQI enzyme, 10q26) | | |
|---|---|---|---|---|---|---|---|---|---|---|---|---|---|---|---|---|---|---|---|---|---|---|---|---|---|
| | | | | | | | | Length (kb) | Units | Allele | Length (kb) | Units | Allele | Units | Allele | Read Count | Units | Allele | Read Count | Units | Allele | Read Count | Units | Allele | Read Count |
| Patient cohort 1 | P01 | F | 37 | 28 | + | 104 | mild myopathic change | ~20 | ~4 | 4qA | ~38.4 | ~10 | 10qA | 4 | 4qA | 6 | 11 | 10qA | 8 | 4.3±0.3 | 4qA | 38 | 11.2±0.1 | - | 19 |
| | | | | | | | | >38 | >10 | 4qB | >63.5 | >17 | 10qA | 22 | 4qB | 5 | 31 | 10qA | 11 | 22.4±0.2 | 4qB | 23 | 31.9±0.3 | - | 10 |
| | P02 | F | 18 | 12 | + | 379 | myopathic change | ~18 | ~3 | 4qA | ~38.4 | ~10 | 10qA | 3 | 4qA | 25 | 11 | 10qA | 45 | - | - | - | - | - | - |
| | | | | | | | | ~98 | ~28 | 4qA | >63.5 | >17 | 10qA | 28 | 4qA | 10 | 25 | 10qA | 24 | | | | | | |
| | P03 | F | 43 | 30 | + | 110 | mild myopathic change | ~18 | ~3 | 4qA | ~38.4 | ~10 | 10qA | 3 | 4qA | 27 | 11 | 10qA | 42 | - | - | - | - | - | - |
| | | | | | | | | >38 | >10 | 4qA | >63.5 | >17 | 10qA | 11 | 4qA | 34 | 29 | 10qA | 20 | | | | | | |
| | P04 | M | 27 | 11 | - | 1406 | myopathic change | ~16 | ~3 | 4qA | ~33.5 | ~8 | 10qA | 3 | 4qA | 18 | 7 | 10qA | 73 | | | | | | |
| | | | | | | | | ~61 | ~17 | 4qB | >63.5 | >17 | 10qA | 19 | 4qB | 42 | 26 | 10qA | 54 | - | - | - | - | - | - |
| | | | | | | | | ~76 | ~21 | 4qA | - | - | - | 23 | 4qA | 30 | - | - | - | | | | | | |
| | P05 | F | 41 | 31 | + | 95 | normal | ~12 | ~2 | 4qA | ~29.9 | ~7 | 10qA | 2 | 4qA | 12 | 7 | 10qA | 57 | | | | | | |
| | | | | | | | | ~38 | ~10 | 4qA | - | - | - | 11 | 4qA | 18 | 18 | 10qA | 30 | - | - | - | - | - | - |
| | | | | | | | | >38 | >10 | 4qA | - | - | - | 17 | 4qA | 60 | | | | | | | | | |
| Patient cohort 2 | P06 | F | 14 | 6 | - | 406 | myopathic change | | | | | | | 2 | 4qA | 3 | 36 | 10qB | 6 | 2.2±0.3 | 4qA | 32 | 34.8±0.2 | - | 14 |
| | | | | | | | | - | - | - | - | - | - | 15 | 4qB | 4 | 48 | 10qA | 6 | 15.0±0.2 | 4qB | 36 | 48.2±0.4 | - | 6 |
| | | | | | | | | | | | | | | 27 | 4qA | 3 | - | - | - | 27.2±0.4 | 4qA | 5 | - | - | - |
| | P07 | M | 23 | 18 | - | 871 | myopathic change | - | - | - | - | - | - | 4 | 4qA | 6 | 7 | 10qA | 12 | - | - | - | - | - | - |
| | | | | | | | | | | | | | | 20 | 4qA | 11 | - | - | - | | | | | | |
| | P08 | M | 18 | 13 | - | 538 | myopathic change | ~21.5 | ~5 | 4qA | ~24.8 | ~6 | 10qA | 5 | 4qA | 6 | 7 | 10qA | 12 | - | - | - | - | - | - |
| | | | | | | | | >63.5 | >17 | 4qA | ~33.5 | ~9 | 10qA | 25 | 4qA | 9 | 10 | 10qA | 8 | | | | | | |
| | P09 | F | 33 | 19 | - | 269 | | ~21.5 | ~5 | 4qA | ~63.5 | ~17 | 10qA | 5 | 4qA | 33 | 16 | 10qA | 33 | - | - | - | - | - | - |

| | | | | | | | | | | | | | | | | | | | | | | | | | |
|---|---|---|---|---|---|---|---|---|---|---|---|---|---|---|---|---|---|---|---|---|---|---|---|---|---|
| | | | | | | | mild myopathic change | >63.5 | >17 | 4qA | | | | 18 | 4qA | 26 | 27 | 10qB | 24 | | | | | | |
| | P10 | F | 39 | 28 | - | 176 | myopathic change | ~15 | ~3 | 4qA | ~24.8 | ~6 | 10qA | 3 | 4qA | 17 | 7 | 10qA | 17 | - | - | - | - | - | - |
| | | | | | | | | >63.5 | >17 | 4qA | >63.5 | >17 | 10qA | 28 | 4qA | 8 | 28 | 10qA | 14 | | | | | | |
| | P11 | M | 20 | 14 | + | 379 | myopathic change | ~15 | ~3 | 4qA | ~55 | ~15 | 10qA | 3 | 4qA | 16 | 14 | 10qA | 18 | - | - | - | - | - | - |
| | | | | | | | | >63.5 | >17 | 4qA | ~63.5 | ~17 | 10qA | 32 | 4qA | 12 | 17 | 10qA | 20 | | | | | | |
| | P12 | F | 15 | 10 | - | 514 | no data | ~12 | ~2 | 4qA | >63.5 | >17 | 10qA | 2 | 4qA | 10 | 21 | 10qB | 13 | - | - | - | - | - | - |
| | | | | | | | | ~48.5 | 15 | 4qA | >63.5 | >17 | 10qA | 16 | 4qA | 18 | 38 | 10qA | 13 | | | | | | |
| | P13 | F | 53 | 32 | + | 341 | myopathic change | ~18.5 | ~4 | 4qA | ~44 | ~12 | 10qA | 4 | 4qA | 18 | 13 | 10qA | 25 | - | - | - | - | - | - |
| | | | | | | | | >63.5 | >17 | 4qB | >63.5 | >17 | 10qA | 18 | 4qB | 18 | 26 | 10qA | 20 | | | | | | |
| Control | C01 | F | - | - | - | - | - | - | - | - | - | - | - | 19 | 4qB | 8 | 15 | 10qA | 21 | 18.8±0.3 | 4qB | 22 | 15.1±0.2 | - | 11 |
| | | | | | | | | | | | | | | 47 | 4qA | 10 | 20 | 10qB | 7 | 46.5±0.5 | 4qA | 13 | 19.9±0.2 | | 9 |
| | C02 | M | - | - | - | - | - | - | - | - | - | - | - | 18 | 4qA | 8 | 17 | 10qB | 8 | 18.6±0.3 | 4qA | 7 | - | - | - |
| | | | | | | | | | | | | | | 20 | 4qA | 9 | 38 | 10qA | 5 | 20.5±0.1 | 4qA | 8 | 38.3±0.1 | | 6 |
| | C03 | M | - | - | - | - | - | - | - | - | - | - | - | 13 | 4qB | 4 | 25 | 10qA | 8 | 12.7±0.5 | 4qB | 20 | 24.9±0.3 | - | 16 |
| | | | | | | | | | | | | | | 22 | 4qB | 4 | 40 | 10qB | 7 | 22.2±0.4 | 4qB | 22 | 40.2±0.5 | | 5 |

**Supplementary Table 3. The relationship between coverage and read length for all samples assayed in the current study.**

| ID | Optical Mapping (Nb.BssSI enzyme) Depth of Coverage(X) | | | | | | | Optical Mapping (Nt.BspQI enzyme) Depth of Coverage(X) | | | | | | |
|---|---|---|---|---|---|---|---|---|---|---|---|---|---|---|
| | >50kb | >100kb | >150kb | >200kb | >300kb | >400kb | >500kb | >50kb | >100kb | >150kb | >200kb | >300kb | >400kb | >500kb |
| P01 | 91.933 | 72.283 | 44.317 | 25.053 | 8.072 | 2.929 | 1.193 | 89.317 | 88.158 | 83.196 | 74.470 | 52.227 | 33.809 | 21.381 |
| P02 | 137.643 | 130.867 | 115.112 | 96.543 | 64.396 | 41.085 | 25.809 | | | | | | | |
| P03 | 131.600 | 127.653 | 116.684 | 99.818 | 62.911 | 35.361 | 19.152 | | | | | | | |
| P04 | 143.400 | 131.871 | 102.360 | 68.998 | 26.527 | 10.404 | 4.460 | | | | | | | |
| P05 | 103.211 | 95.312 | 77.234 | 57.162 | 29.157 | 14.876 | 7.880 | | | | | | | |
| P06 | 78.593 | 75.406 | 67.226 | 57.073 | 37.747 | 23.331 | 14.030 | 102.041 | 100.567 | 94.055 | 82.001 | 51.249 | 28.595 | 15.339 |
| P07 | 48.838 | 48.838 | 48.838 | 42.520 | 28.053 | 17.199 | 10.296 | | | | | | | |
| P08 | 39.261 | 32.718 | 19.751 | 11.085 | 5.285 | 2.710 | 1.254 | | | | | | | |
| P09 | 124.255 | 120.606 | 108.099 | 87.964 | 45.245 | 18.203 | 6.587 | | | | | | | |
| P10 | 99.546 | 93.879 | 76.014 | 52.752 | 18.833 | 5.301 | 1.370 | | | | | | | |
| P11 | 127.016 | 124.958 | 118.272 | 106.309 | 71.998 | 41.007 | 22.011 | | | | | | | |
| P12 | 128.750 | 124.503 | 112.920 | 95.891 | 59.181 | 32.112 | 16.769 | | | | | | | |
| P13 | 90.945 | 87.330 | 77.280 | 63.387 | 36.743 | 19.762 | 10.407 | | | | | | | |
| C01 | 62.014 | 61.364 | 59.068 | 55.277 | 45.117 | 34.434 | 25.572 | 87.706 | 86.220 | 81.378 | 74.310 | 57.787 | 42.060 | 29.659 |
| C02 | 158.290 | 134.794 | 78.313 | 33.574 | 7.460 | 2.786 | 1.140 | 75.913 | 75.867 | 66.128 | 52.934 | 28.986 | 14.820 | 7.350 |
| C03 | 40.098 | 39.174 | 34.342 | 27.950 | 16.434 | 8.696 | 4.280 | 120.868 | 117.484 | 106.773 | 92.367 | 64.655 | 44.068 | 29.639 |
| GM19238 | 87.668 | 84.228 | 72.936 | 57.914 | 33.187 | 18.884 | 10.885 | 134.113 | 133.117 | 127.506 | 117.539 | 91.381 | 66.599 | 47.822 |
| GM19239 | 56.751 | 54.657 | 47.554 | 37.024 | 16.961 | 6.085 | 1.951 | 82.644 | 81.800 | 77.206 | 70.043 | 52.995 | 37.104 | 25.397 |
| GM19240 | 83.675 | 81.083 | 72.446 | 59.581 | 33.530 | 16.094 | 7.128 | 68.590 | 67.690 | 63.515 | 56.823 | 40.398 | 26.065 | 16.129 |
| HG00512 | 59.304 | 57.548 | 51.376 | 42.244 | 24.333 | 12.161 | 5.590 | 59.261 | 58.028 | 53.592 | 47.526 | 34.890 | 24.654 | 17.491 |
| HG00513 | 79.074 | 75.680 | 64.498 | 49.726 | 25.667 | 12.811 | 6.526 | 72.632 | 71.708 | 66.956 | 59.306 | 41.423 | 26.921 | 17.183 |
| HG00514 | 98.434 | 95.986 | 87.993 | 76.215 | 50.440 | 29.631 | 16.112 | 70.860 | 70.061 | 66.064 | 59.786 | 44.815 | 31.517 | 21.888 |
| HG00731 | 63.378 | 61.456 | 54.091 | 42.770 | 21.785 | 9.799 | 4.212 | 53.520 | 52.547 | 47.738 | 39.825 | 23.189 | 12.136 | 6.481 |
| HG00732 | 60.728 | 58.987 | 53.176 | 44.699 | 27.181 | 14.688 | 7.420 | 170.568 | 147.766 | 104.465 | 76.238 | 43.652 | 25.518 | 15.170 |

| | | | | | | | | | | | | | |
|---|---|---|---|---|---|---|---|---|---|---|---|---|---|
| HG00733 | 101.372 | 97.054 | 83.851 | 66.465 | 35.931 | 17.050 | 7.576 | 95.759 | 94.104 | 86.952 | 76.720 | 54.472 | 35.963 | 23.059 |

**Supplementary Table 4. In silico analysis of the position of D4Z4 repeat units on 4q35 in the human reference genome GRCh38 (note that a previous study incorrectly determined repeat units in GRCh38). We used blastn to map D38024.1 to reference sequence and obtained position information. We failed to detect Nt.BspQI labels in repeat unit #2 and #5 in data sets that we have analyzed in the current study, suggesting that GRCh38 may have included a very rare allele or may contain assembly errors in this region.**

| Unit | Chr | Start | End | Length | Nt.BspQI recognition site |
|---|---|---|---|---|---|
| **1** | 4 | 190066141 | 190069434 | 3294 | - |
| **2** | 4 | 190069435 | 190072731 | 3297 | 190070830 |
| **3** | 4 | 190072732 | 190076024 | 3293 | - |
| **4** | 4 | 190076025 | 190079317 | 3293 | - |
| **5** | 4 | 190079318 | 190082611 | 3294 | 190080713 |
| **6** | 4 | 190082612 | 190085911 | 3300 | - |
| **7** | 4 | 190085912 | 190089204 | 3293 | - |
| **8** | 4 | 190089205 | 190092504 | 3300 | - |

**Supplementary Table 5. In silico analysis of the position of D4Z4 repeat units on 10q26 from GRCh38. We used blastn to map D38024.1 to reference sequence and obtained position information. The presence of a gap between unit 1-6 and unit 7-12 is due to genome assembly errors introduced in GRCh38, but it is not present in GRCh37.**

| Unit | chr | start | end | length |
|---|---|---|---|---|
| **1** | 10 | 133665335 | 133668644 | 3310 |
| **2** | 10 | 133668645 | 133671943 | 3299 |
| **3** | 10 | 133671944 | 133675242 | 3299 |
| **4** | 10 | 133675243 | 133678552 | 3310 |
| **5** | 10 | 133678553 | 133681862 | 3310 |
| **6** | 10 | 133681863 | 133685161 | 3299 |
| **Gap** | | | | |
| **7** | 10 | 133741514 | 133744823 | 3310 |
| **8** | 10 | 133744824 | 133748133 | 3310 |
| **9** | 10 | 133748134 | 133751442 | 3309 |
| **10** | 10 | 133751443 | 133754741 | 3299 |
| **11** | 10 | 133754742 | 133758051 | 3310 |
| **12** | 10 | 133758052 | 133761350 | 3299 |

**Supplementary Table 6. In silico analysis of the position of D4Z4 repeat units on KQ983257.1 from GRCh38.p7. We used blastn to map D38024.1 to reference sequence and obtained position information.**

| Unit | Chr | Start | End | Length |
|---|---|---|---|---|
| **1** | KQ983257.1 | 158744 | 162049 | 3306 |
| **2** | KQ983257.1 | 162050 | 165355 | 3306 |
| **3** | KQ983257.1 | 165356 | 168661 | 3306 |
| **4** | KQ983257.1 | 168662 | 171967 | 3306 |
| **5** | KQ983257.1 | 171968 | 175273 | 3306 |
| **6** | KQ983257.1 | 175274 | 178579 | 3306 |
| **7** | KQ983257.1 | 178580 | 181885 | 3306 |
| **8** | KQ983257.1 | 181886 | 185191 | 3306 |
| **9** | KQ983257.1 | 185192 | 188497 | 3306 |
| **10** | KQ983257.1 | 188498 | 191803 | 3306 |
| **11** | KQ983257.1 | 191804 | 195109 | 3306 |
| **12** | KQ983257.1 | 195110 | 198415 | 3306 |
| **13** | KQ983257.1 | 198416 | 201721 | 3306 |

### Supplementary Figures

**Supplementary Figure 1. Determination of D4Z4 copy number on individuals affected with FSHD and control subjects.** Screenshots from the Bionano Access browser are shown, and for each panel, the labeling enzyme is annotated. For repeat quantification using the Nt.BspQI enzyme, mean ± SD (standard deviation) is annotated in the figure.

P01(4q35)

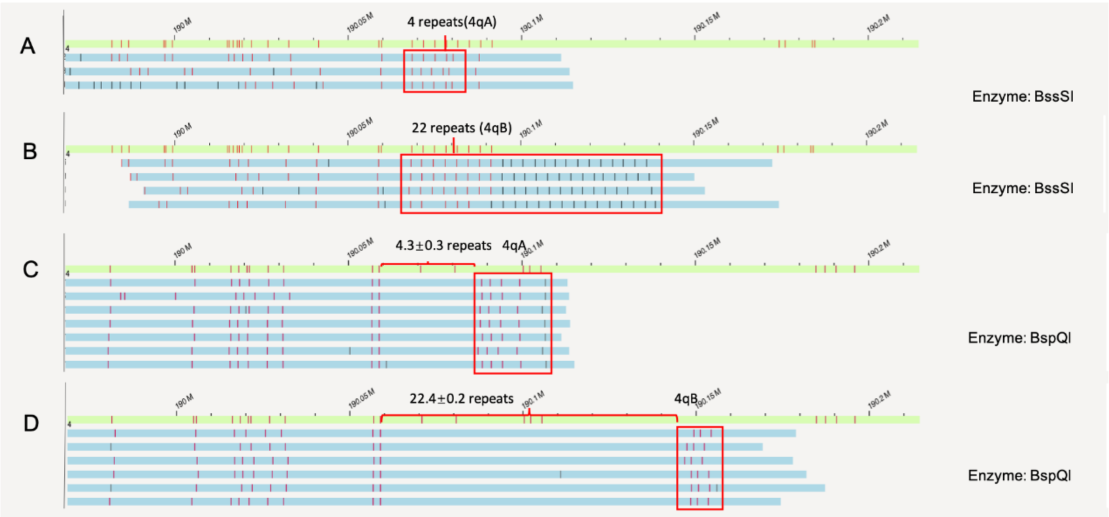


P01(10q26)

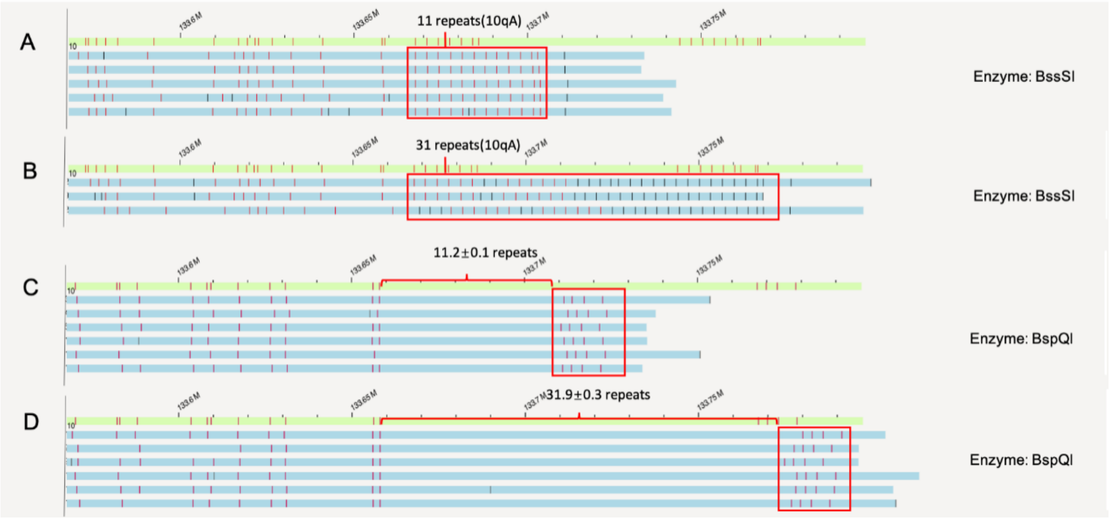

# P02(4q35)

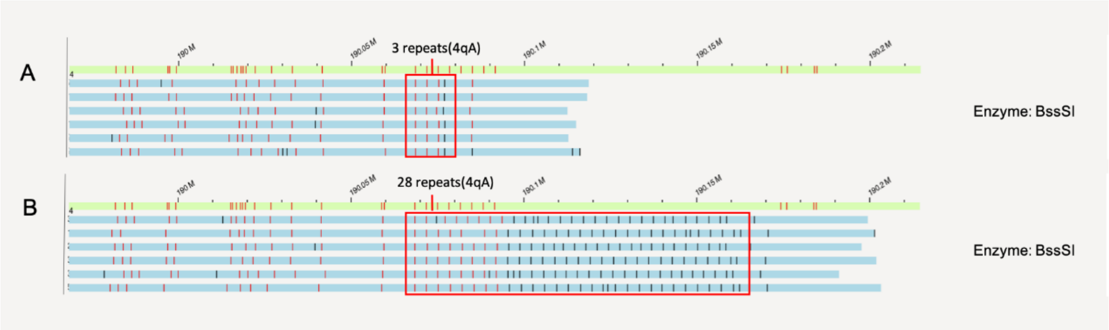


# P02(10q26)

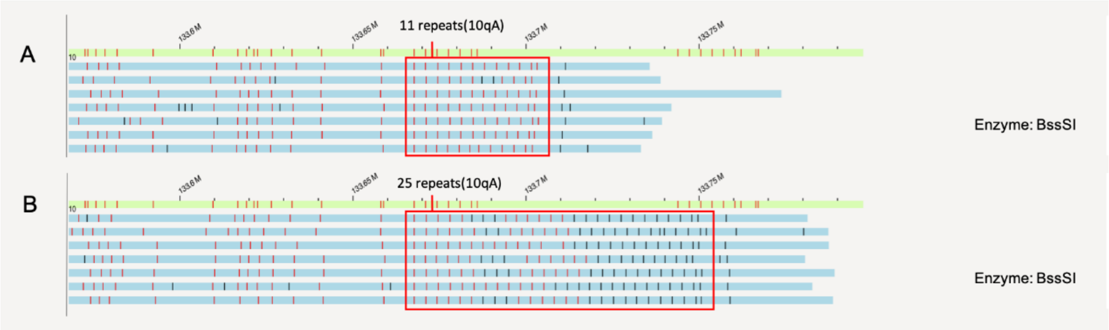


# P03(4q35)

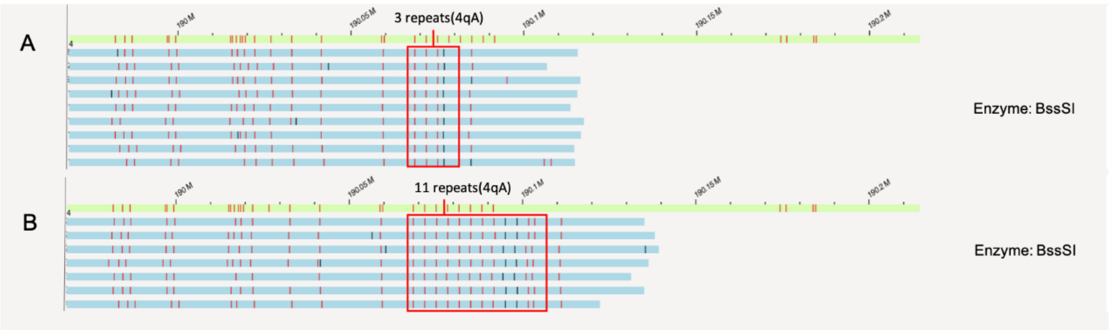


# P03(10q26)

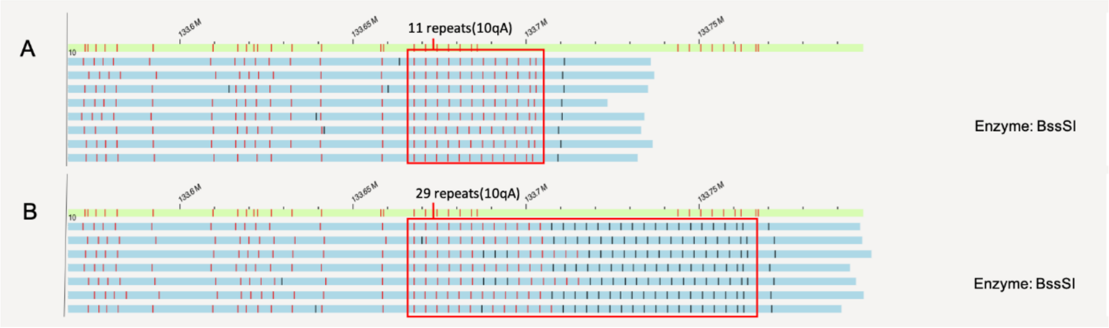

# P04(4q35)

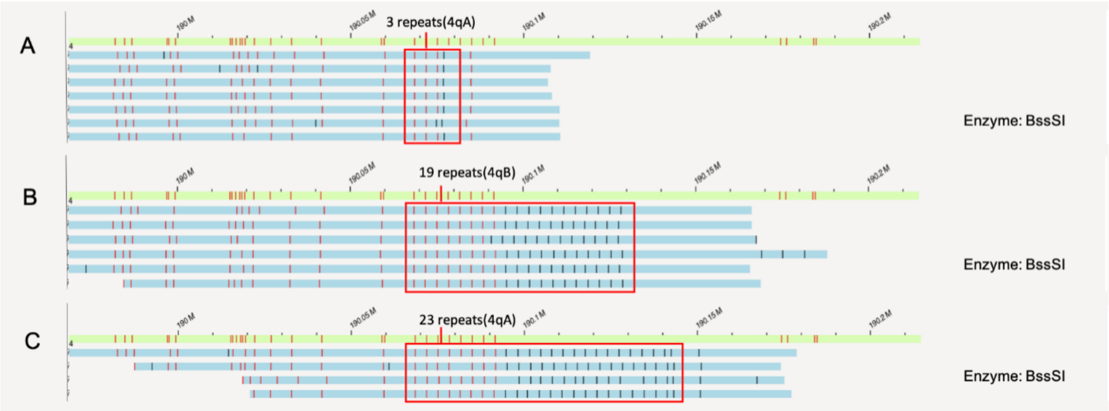


# P04(10q26)

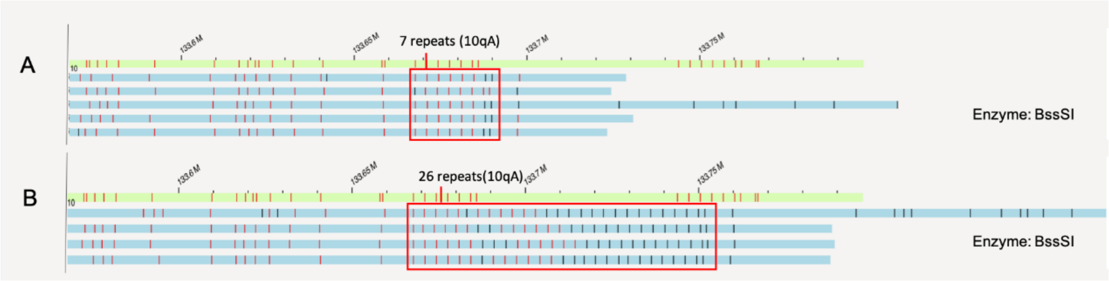


# P05(4q35)

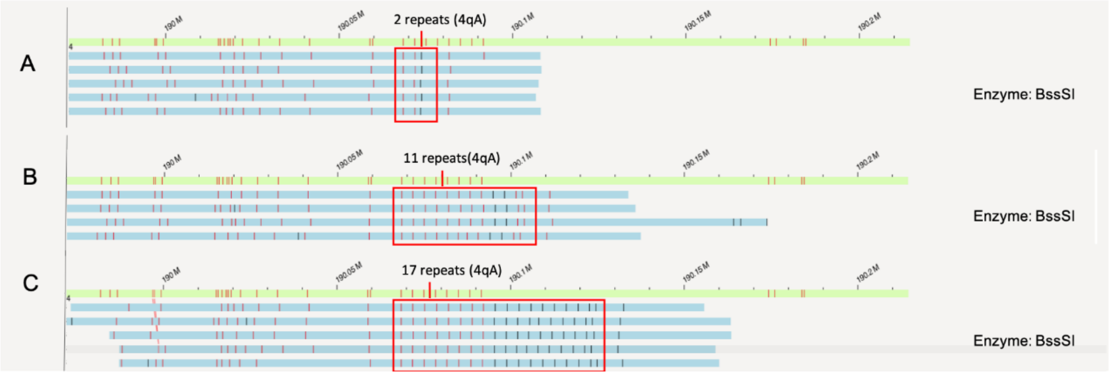


# P05(10q26)

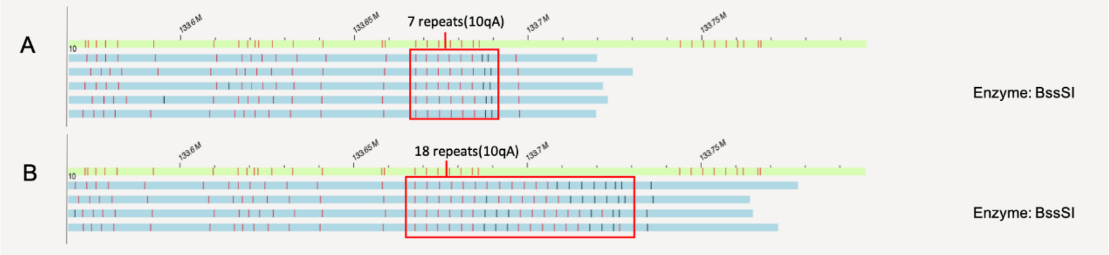

# P06(4q35)

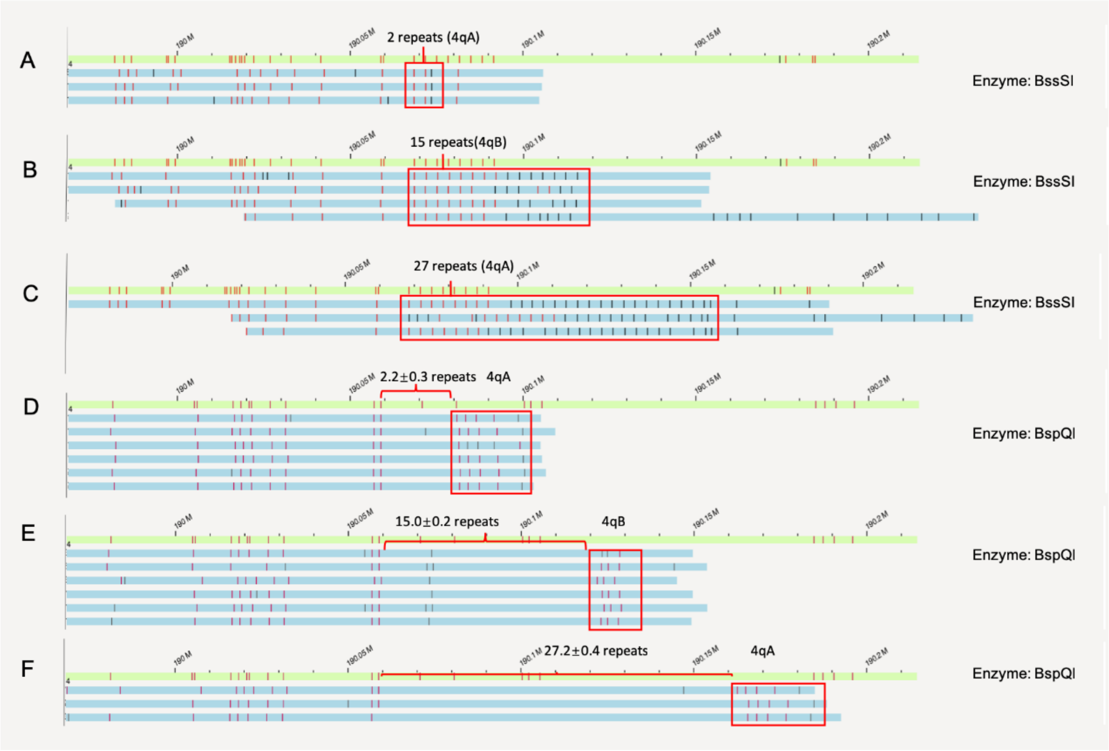


# P06(10q26)

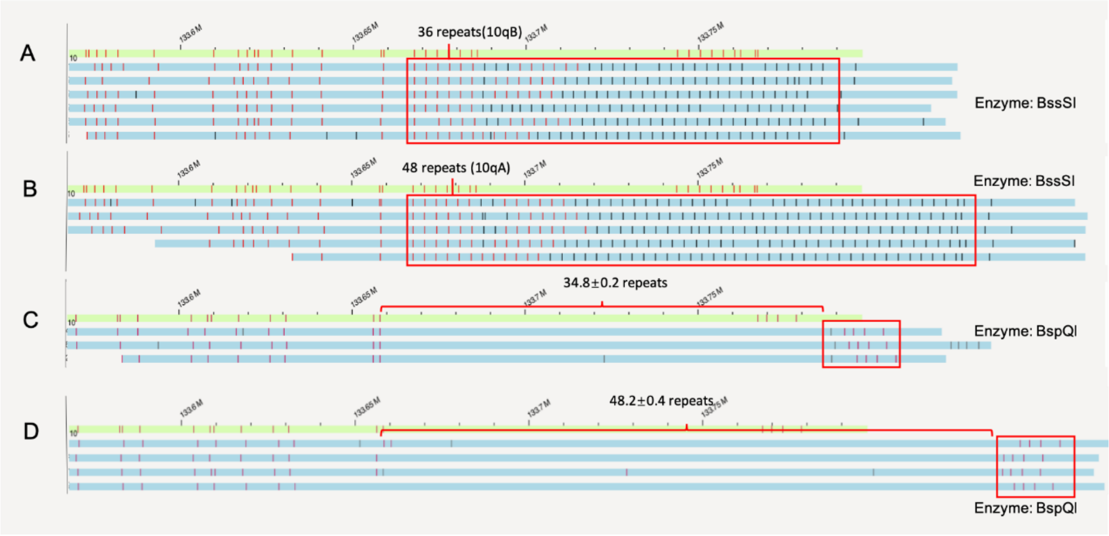


# P07(4q35)

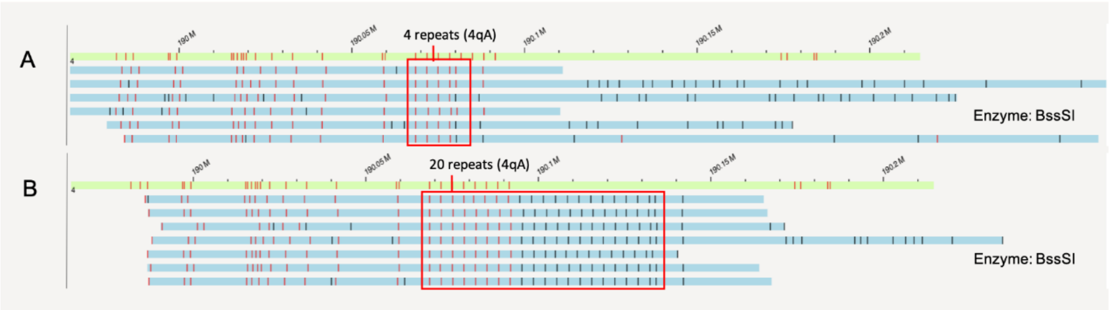

# P07(10q26)

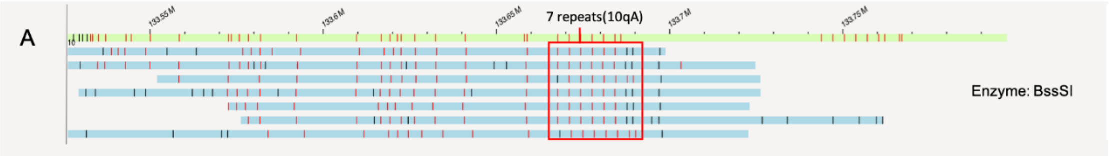


# P08(4q35)

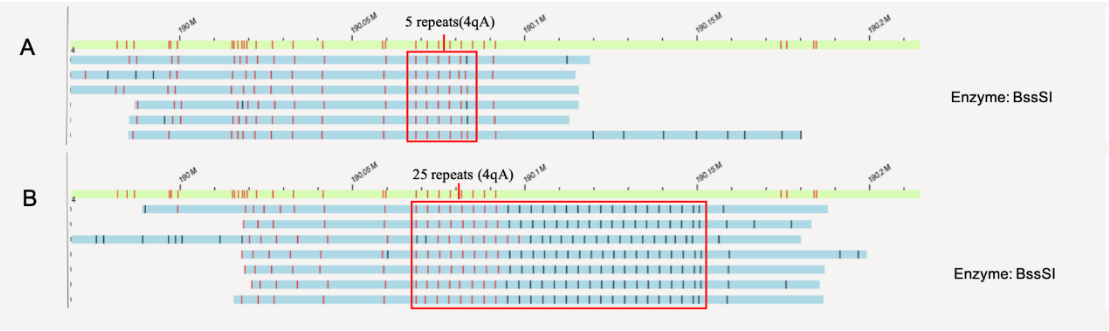


# P08(10q26)

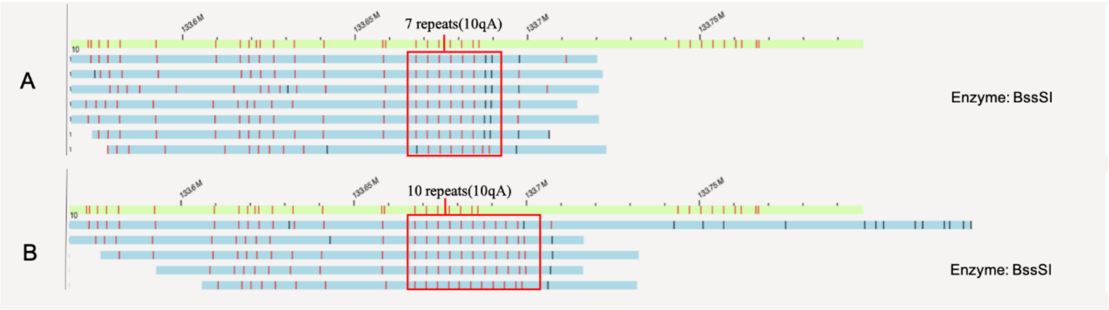


# P09(4q35)

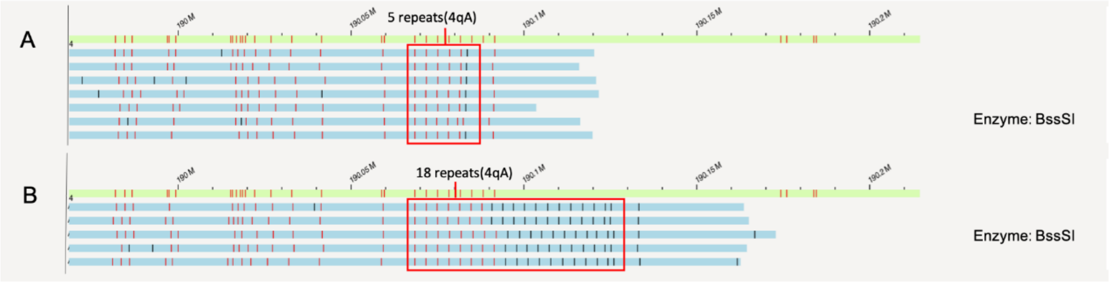


# P09(10q26)

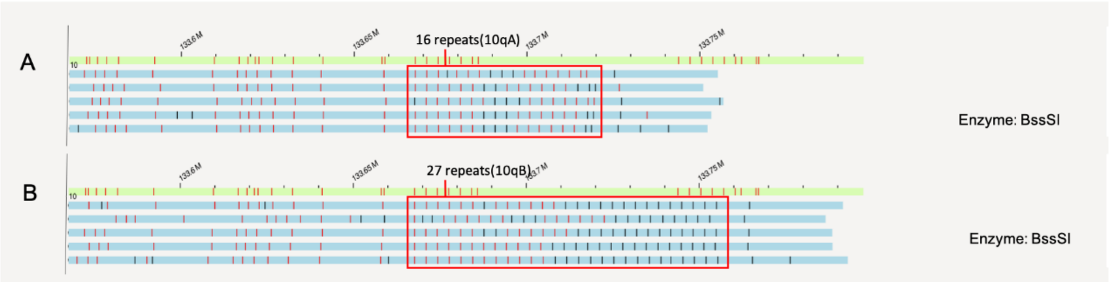

## P10(4q35)

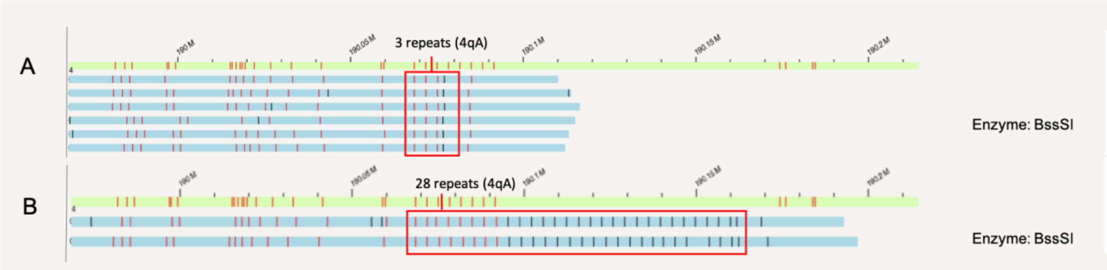


## P10(10q26)

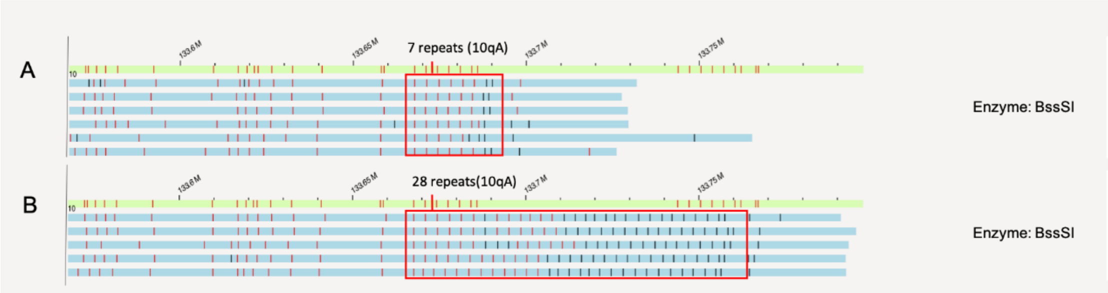


## P11(4q35)

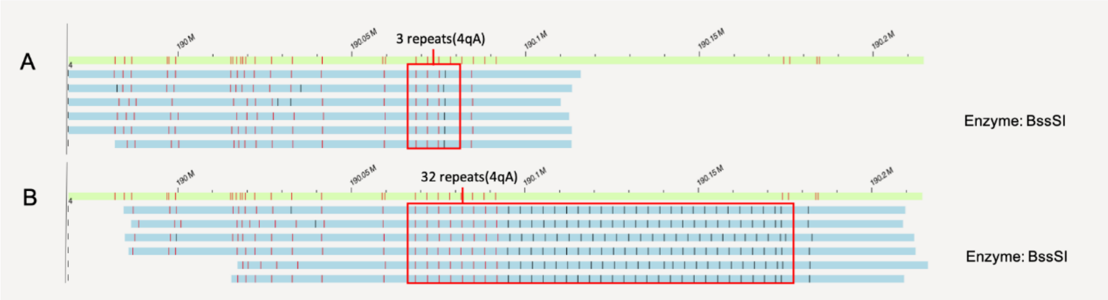


## P11(10q26)

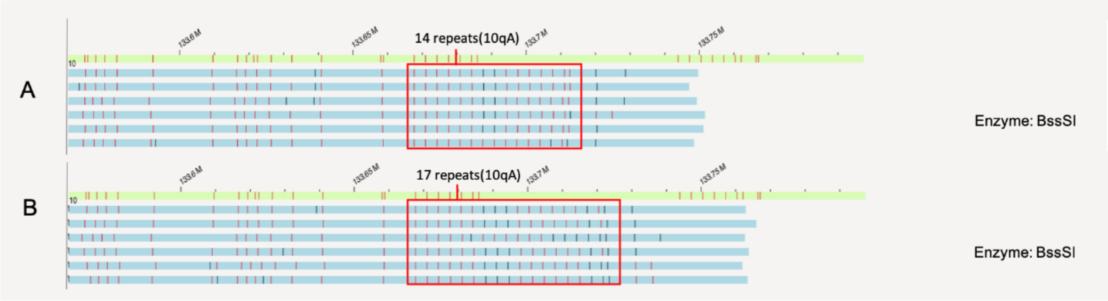

# P12(4q35)

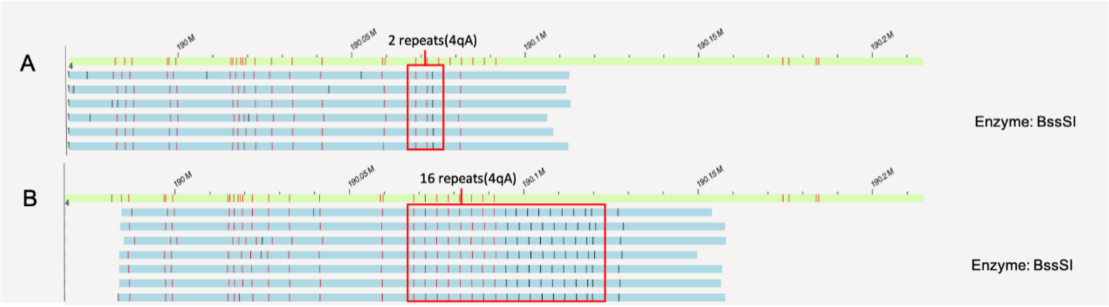


# P12(10q26)

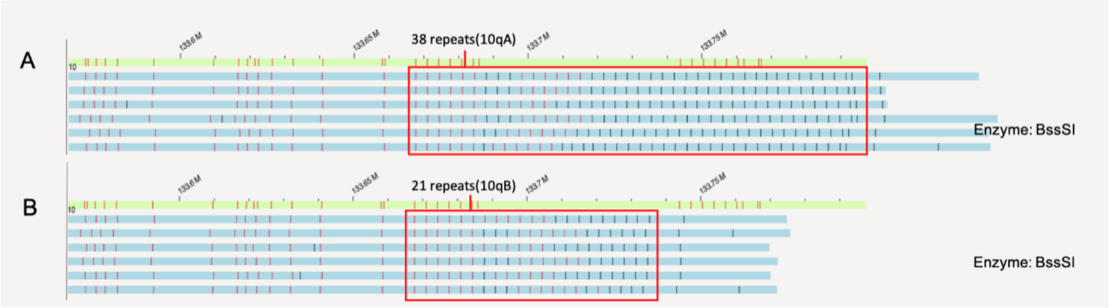


# P13(4q35)

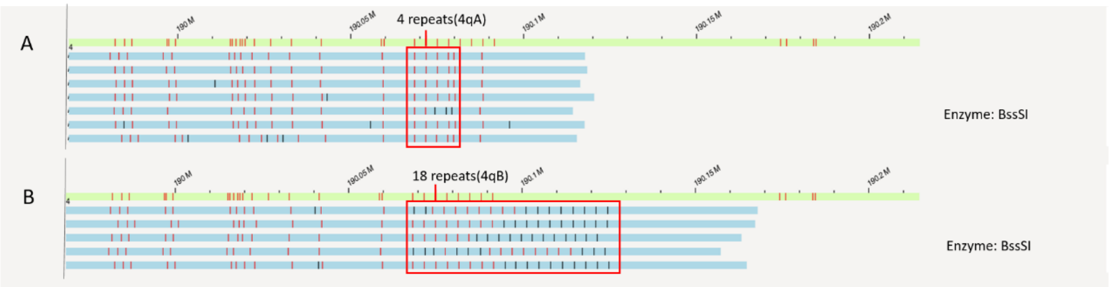


# P13(10q26)

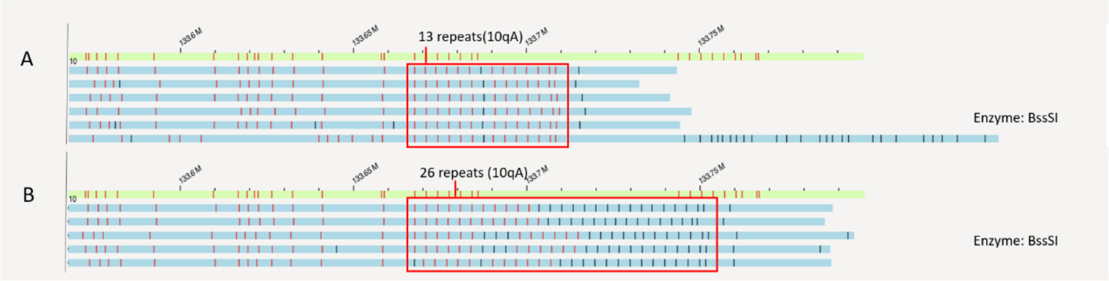

# C01(4q35)

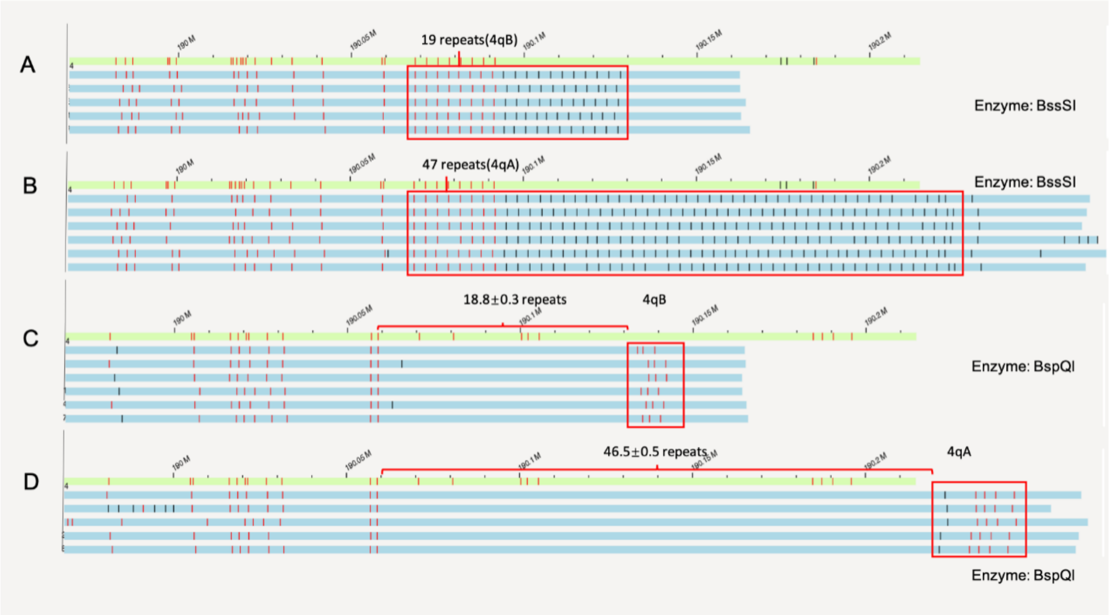


# C01(10q26)

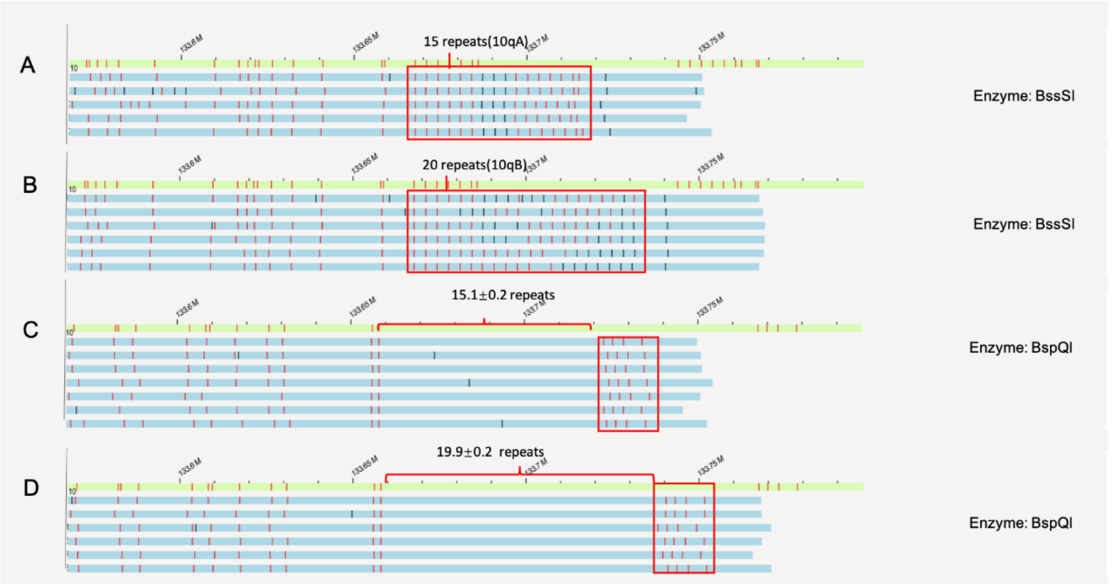

# C02(4q35)

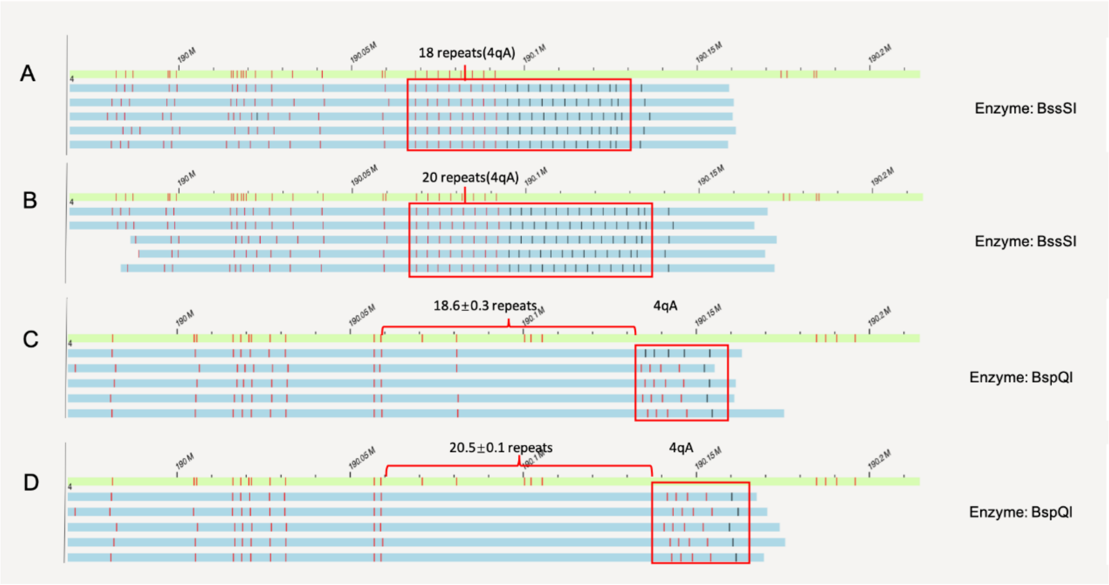


# C02(10q26)

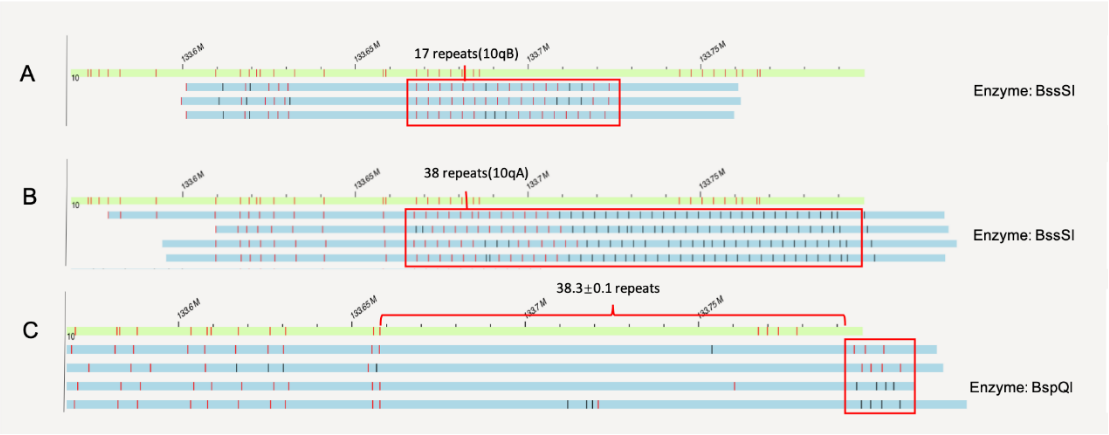

# C03(4q35)

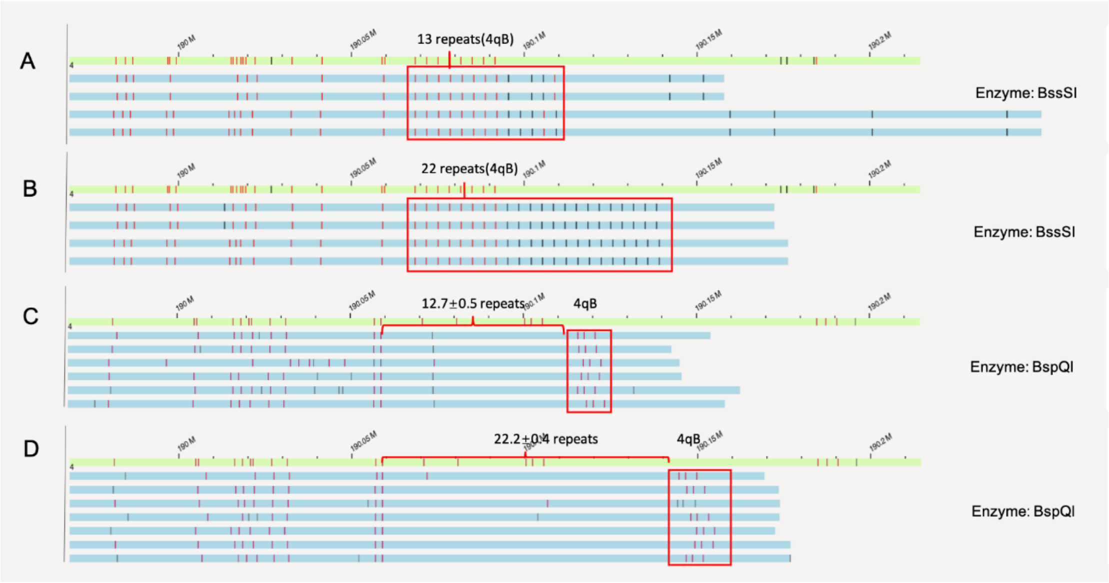


# C03(10q26)

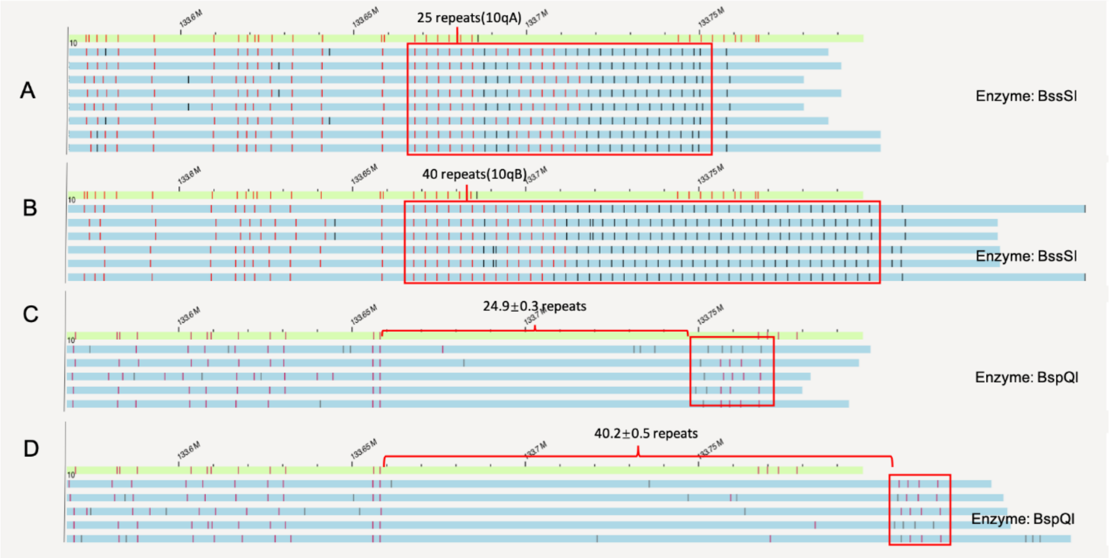

HG00512

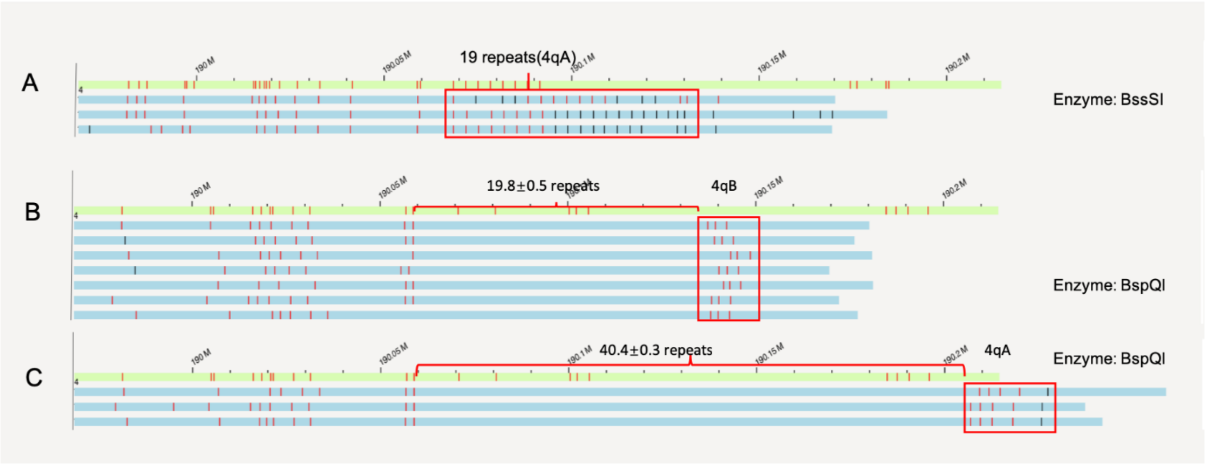


HG00513

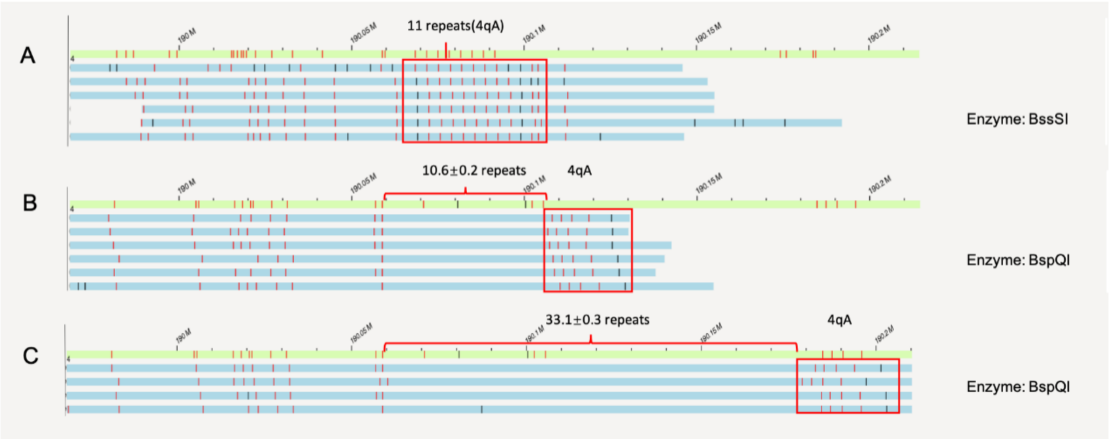


HG00514

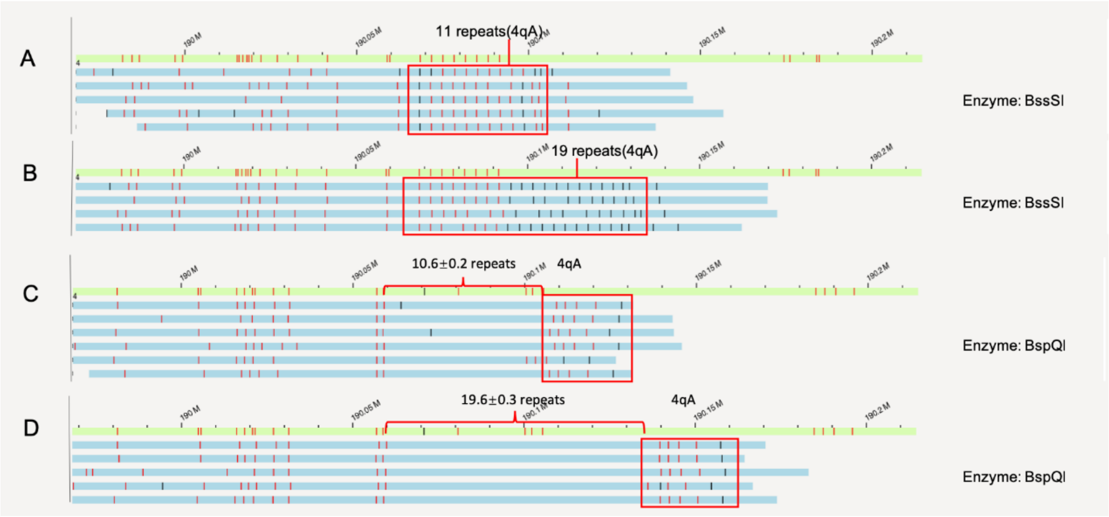

# HG00731

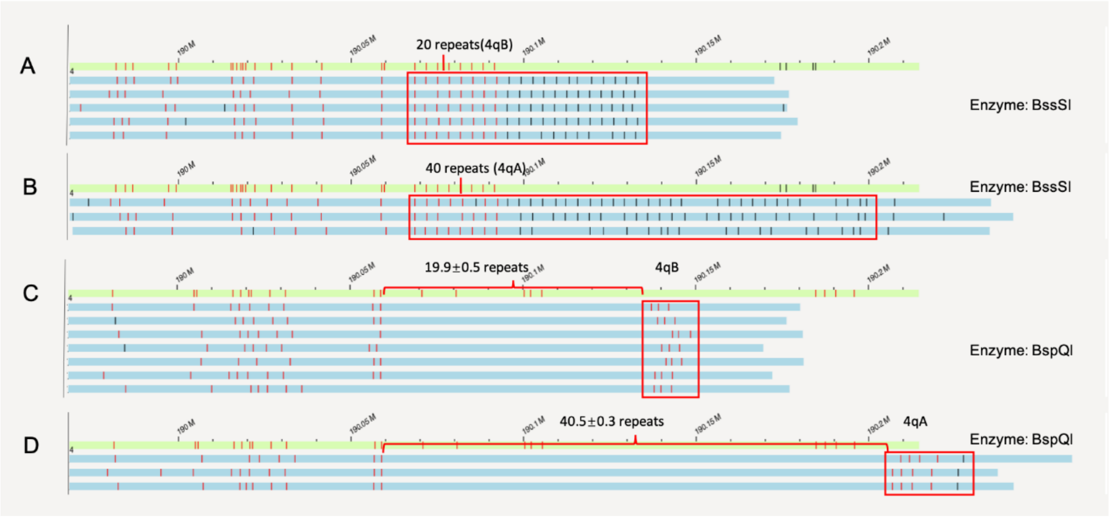


# HG00732

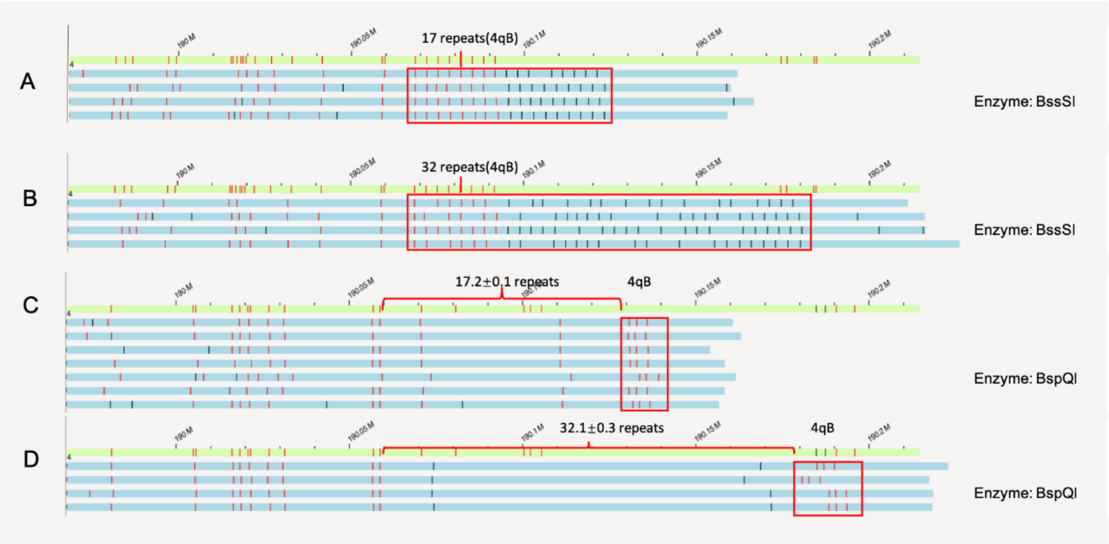

# HG00733

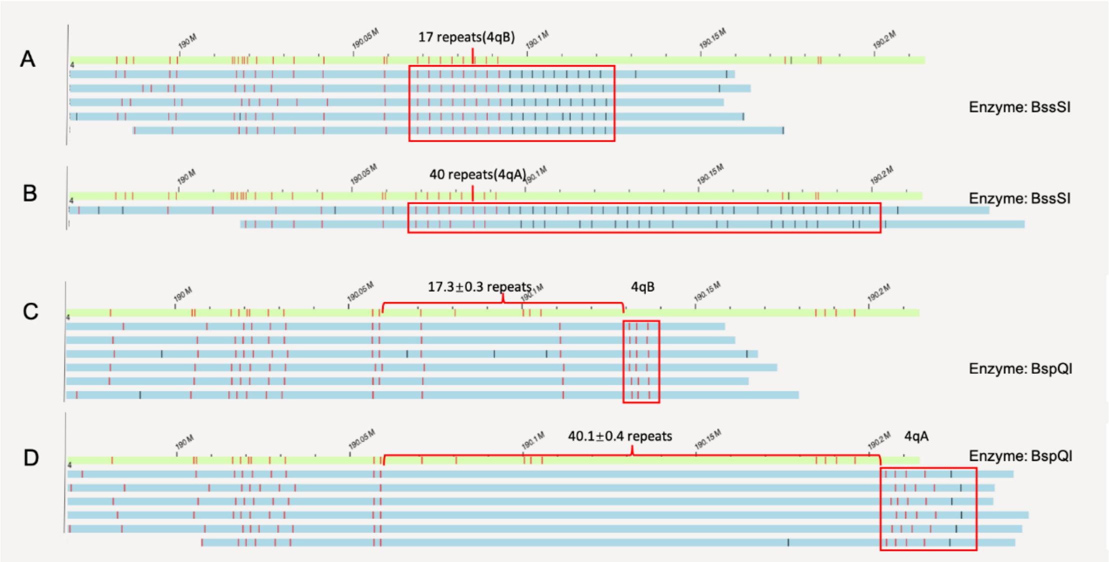


# GM19238

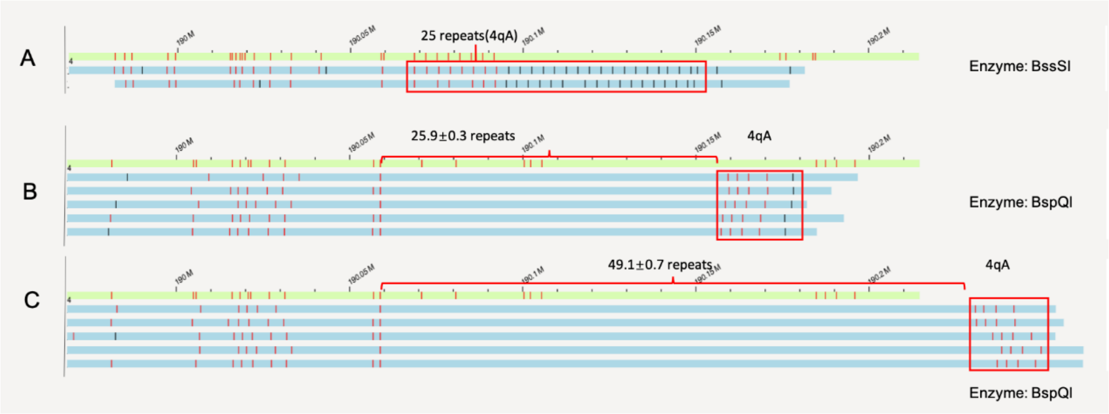


# GM19239

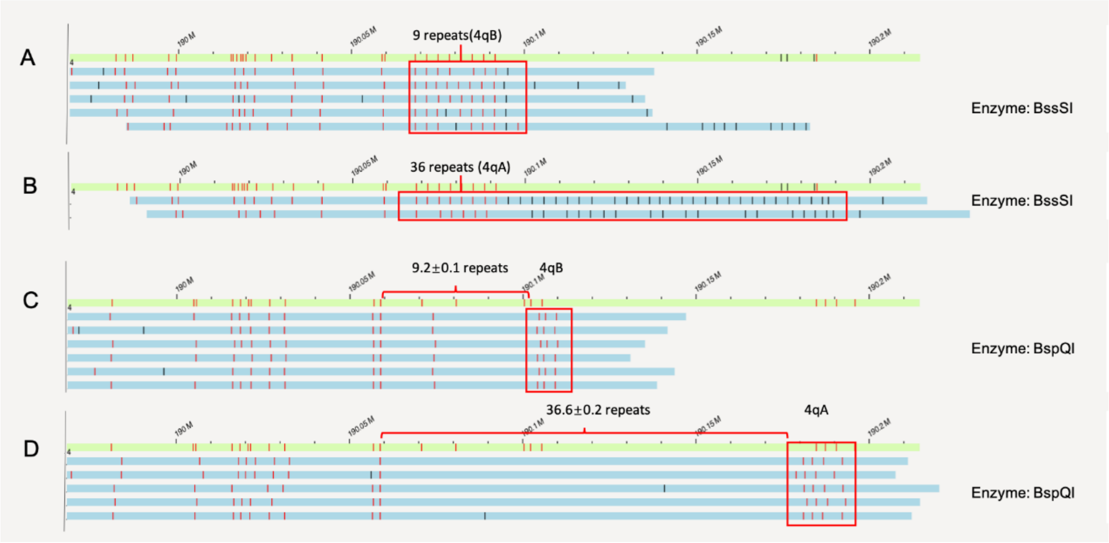

# GM19240

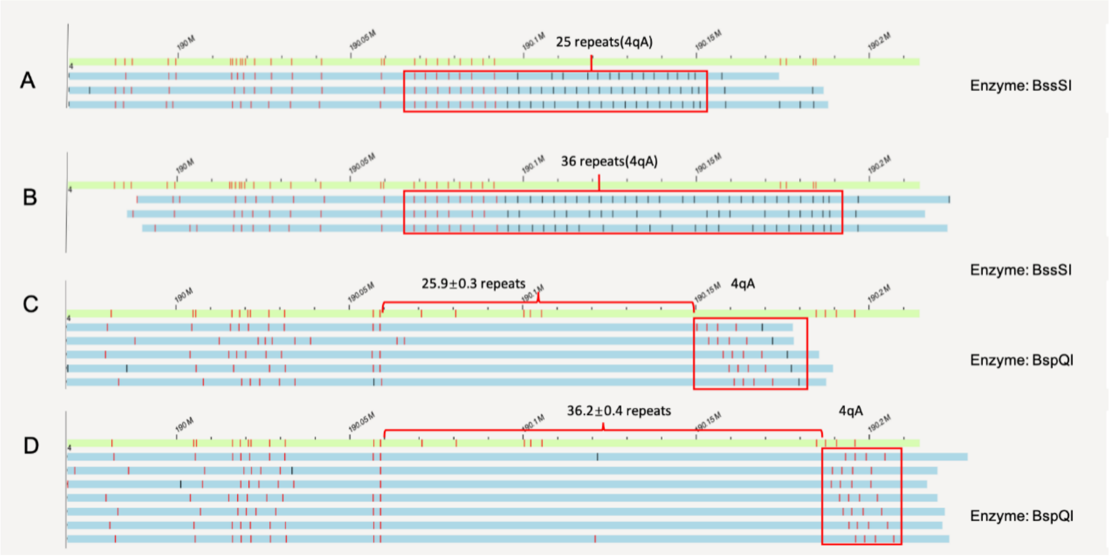
A
25 repeats(4qA)
Enzyme: BssSI
B
36 repeats(4qA)
Enzyme: BssSI
C
25.9±0.3 repeats
4qA
Enzyme: BspQI
D
36.2±0.4 repeats
4qA
Enzyme: BspQI

**Supplementary Figure 2. Southern blots on patients with FSHD from cohort 2.** DNA samples for P06/P07 were not available for Southern blot. E/H and p13E-11: Double digested with EcoRI/HindIII and then labeled with probe p13E-11, and all the 4q and 10q segments are illustrated. E/B and p13E-11: Double digested with EcoRI/BlnI and then labeled with probe p13E-11, and the 10q segments are digested so only 4q segments are illustrated. H and 4qA: digested with HindIII and then labeled with probe 4qA, and the 4qA alleles are illustrated. H and 4qB: digested with HindIII and then labeled with probe 4qB, and the 4qB alleles are illustrated. The star "*" denotes pathogenic allele with <10 repeat units and with a 4qA configuration. The plus sign "+" denotes somatic mosaic allele.

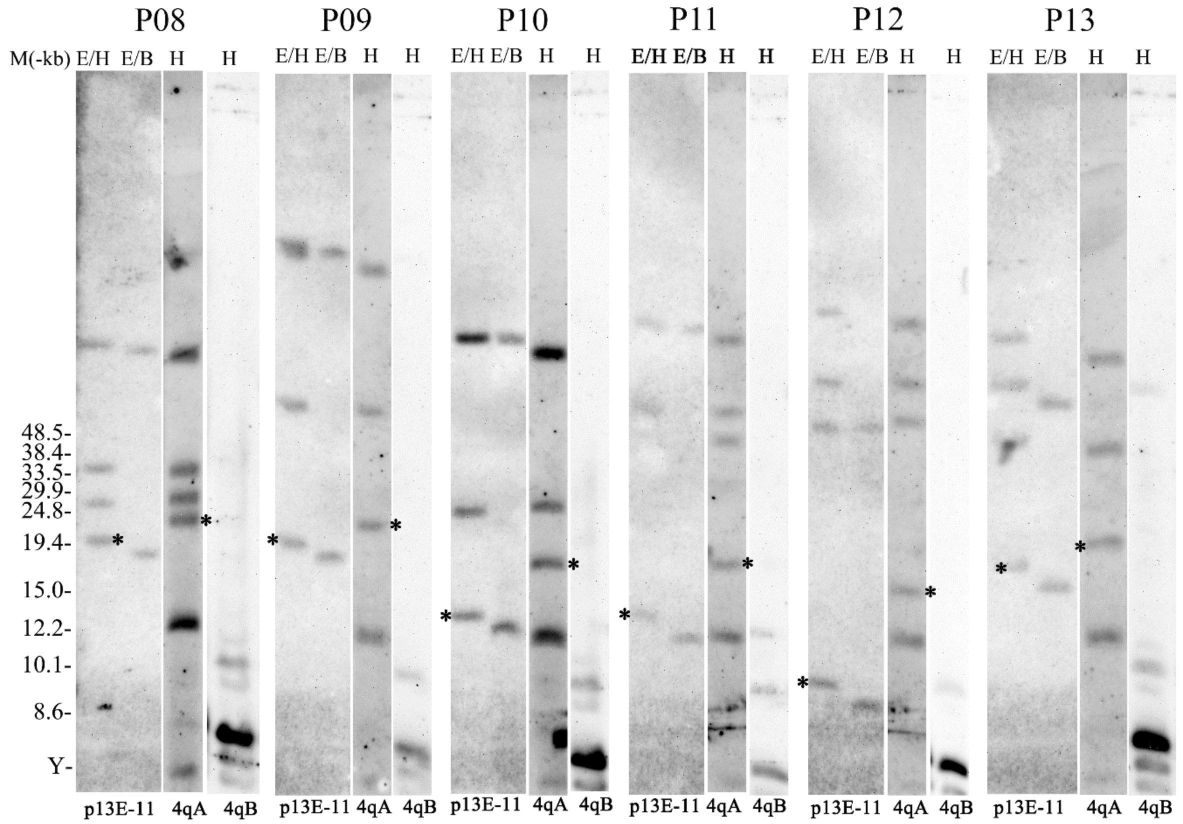
